# Multiomics analysis identifies VPA-induced changes in neural progenitor cells, ventricular-like regions, and cellular microenvironment in dorsal forebrain organoids

**DOI:** 10.1101/2025.06.24.661272

**Authors:** Zeynep Yentür, Kseniia Sarieva, Lizia Branco, Theresa Kagermeier, Christina Kulka, Mohamed A. Jarboui, Katharina Becker, Simone Mayer

## Abstract

Pharmaceutical agents, such as antiepileptic medications, can cross fetal barriers and affect the developing brain. Prenatal exposure to the antiepileptic drug valproate (VPA) is associated with an increased risk of neurodevelopmental disorders, including congenital malformations and autism spectrum disorder. In animal models and neural organoids, VPA has been shown to alter signaling pathways, such as Wnt pathway, providing insights into VPA-induced neurodevelopmental defects. Here, we exposed dorsal forebrain organoids to VPA for 30 days and examined effects at the tissue, cellular, and molecular level. VPA treatment disrupted ventricular-like regions, indicating defects in cell-cell and cell-matrix interactions. Transcriptomics analysis confirmed altered expression of extracellular matrix (ECM) genes and single cell RNA sequencing analysis identified genes involved in microenvironment sensing, such as cellular mechanosensing and Hippo-YAP/TAZ signaling pathway. Finally, proteomics analysis corroborated that VPA alters the microenvironment of the human dorsal forebrain organoids by disrupting the secretion of ECM proteins. Altogether, our study suggests that VPA-treated dorsal forebrain organoids serve as a model to investigate the role of extracellular processes in brain development and to understand how their disruptions might contribute to neurodevelopmental disorders.

## Introduction

Environmental factors, including maternal medications, have been implicated in shaping early brain development (1). Some antiepileptic drugs (AEDs), such as Levetiracetam (LEV), are considered non-teratogenic and safe for use during pregnancy (2). However, other medications that can pass maternal and fetal barriers might increase the risk of neurodevelopmental and/or psychiatric disorders (1). For example, valproate (VPA), an AED, is a well-known teratogen that passes the barriers and reaches the fetal brain (3-5). *In utero* exposure to VPA, especially in the first trimester, is associated with increased risk of congenital malformations, such as neural tube closure defects, and neurodevelopmental disorders, such as autism spectrum disorder (ASD)(6, 7). Despite its known teratogenic effects, VPA treatment may be continued during pregnancy when it is the only effective treatment for seizure control in patients who do not respond to other AEDs (8). Since the use of VPA is unavoidable in some cases of pregnancies of epileptic patients, understanding the mechanism by which VPA disrupts neurodevelopment is crucial to develop strategies to mitigate the risk.

*In utero* exposure to VPA is a common environmentally induced animal model of autism spectrum disorder (ASD)(9). Rodents prenatally exposed to VPA display ASD-like behavioral phenotypes such as reduced sociability and repetitive hyperactivity combined with reduced exploratory activity (10-13). This animal model also provides a tool to study the neurobiology of VPA-induced ASD and VPA-induced structural malformations in the nervous system. Findings show that VPA results in dysregulated differentiation of neural progenitor cells (14) and excitation/inhibition imbalance (15). Additionally, the expression level of collagens was increased in the medial prefrontal cortex of rats prenatally exposed to VPA (16). A possible mechanism that might induce these changes is through epigenetic effect of VPA as a histone deacetylase inhibitor (HDACi)(17). However, animal models are limited in understanding the VPA-induced ASD pathophysiology and the effects of VPA on the developing human brain due to interspecies differences (18).

Human induced pluripotent stem cell (iPSC)- and embryonic stem cell (ESC)-derived neural organoids provide an *in vitro* model system to investigate the mechanisms of environmental exposures in neurodevelopment (1, 19, 20) and developmental defects in ASD (21, 22) in a human-specific tissue context. Various neural organoid protocols have been developed, including regionalized protocols such as dorsal forebrain organoids (DFOs), which recapitulate important features of the cellular, molecular, and cytoarchitectural features of specific regions of the human brain (23, 24). Human neural organoids have been utilized to study the effects VPA exposure and its potential link to ASD (25-27). Independent studies have shown that VPA treatment altered ventricular zone (VZ)-like structures and progenitors, and impaired differentiation in neural organoids (26-30). While these studies have provided insights into the cellular and transcriptional changes induced by VPA exposure, they have not investigated how microenvironment components, such as extracellular matrix (ECM) are altered in a human tissue context.

In this study, we used human iPSC-derived DFOs to examine the cellular and molecular effects of VPA exposure during early neocortical development. We treated the organoids with VPA from day (D)21 of differentiation until D50 to recapitulate continuous exposure of progenitor cells to VPA at early stages of their expansion and neural differentiation. Using immunohistochemistry, we found that VPA exposure disrupted the VZ-like structures, possibly by disrupting the cellular microenvironment. We identified changes in intracellular pathways related to microenvironment sensing such as mechanosensing via integrin and PIEZO channels and Hippo-YAP/TAZ signaling pathway especially in progenitor cells upon VPA treatment through single-cell transcriptomic analysis. Following up on this finding, we analyzed the secretome using mass spectrometry and found increased secretion of ECM components. In conclusion, for the first time, we show an altered microenvironment in VPA-exposed human neural organoids, opening the door to new therapeutic approaches to circumvent VPA-induced neurodevelopmental defects.

## Materials and Methods

### iPSC culture

Commercially available human induced pluripotent stem cell (iPSC) lines BIONi010-C (Source: EBiSC), KOLF2.1J (Source: The Jackson Laboratory), HMGU1 (Source: Helmholtz Zentrum München Deutsches Forschungszentrum für Gesundheit und Umwelt (GmbH))(31) were used for the experiments. The iPSC lines were cultured in standard conditions at 37°C, 5% CO2 and 100% humidity. The cells were maintained in E8 flex medium (Gibco, Cat. no. A2858501) or mTESR Plus media (STEMCELL Technologies, Cat. no. 100-0276) and passaged onto hESC-qualified growth factor-reduced Matrigel-coated (Corning, Cat. no. 354277) or Geltrex-coated (ThermoFisher Scientific, Cat. no. A1413302) plates in colonies using Gentle Dissociation Reagent (STEMCELL Technologies, Cat. no. 07174). The media after passaging was supplemented with Thiazovivin (Sigma-Aldrich, Cat. no. 420220) and replaced with fresh media the next day. All cell lines were tested for pluripotency with antibodies against OCT4 (rabbit, 1:500, Abcam, Cat. no. ab19857) and mycoplasma contamination (TaKaRa, Cat. no. 6601) upon thawing. Cells with passage number below 25 were used for the experiments.

### Dorsal forebrain organoid differentiation

Dorsal forebrain organoids were generated by using previously described protocol (23) with minor changes. iPSCs that reached around 80% confluency were treated with Accutase (Merck, Cat. no. A6964) to obtain single cell dissociation. Then, 9000 cells per well were seeded into 96 well V-bottom low adhesion plates (S-bio, Cat. no. MS-9096VZ) in cortical differentiation medium (CDM) I (Glasgow’s MEM (Gibco, Cat. no. 11710035) with 20% KnockOut Serum Replacement (KOSR, Thermofisher Scientific, Cat. no. 10828028), 1X non-essential amino acids (Sigma, Cat. no. M7145), 0.11 mg/mL Sodium Pyruvate (Thermofisher Scientific, Cat. no. 11360070), 1X Penicillin-Streptomycin (Thermofisher Scientific, Cat. no. 15140122), 0.1 mM β-Mercaptoethanol (Gibco, Cat. no. 21985023) with supplements (20 µM Y-27632 (Cayman Chemical, Cat. no. 10005583), 5 µM SB 431542 (Tocris, Cat. no. 1614), and 3 µM IWR-1 (Merck, Cat. no. 681669)). Alternatively, 9000 cells per well were seeded in specific iPSC media for each cell line. The next day, the aggregate formation was checked, and the media was replaced with supplemented CDMI as written in the protocol. Media change was performed every three days and on day six Y-27632 was removed from the media. On day 18, organoids were transferred to 24-well low adhesion plates in CDMII (DMEM/F12 + 1X Glutamax supplement (Thermofisher Scientific, Cat. no. 31331093) with 1X N2 Supplement (Thermofisher Scientific, Cat. no. 17502048), 1X CD Lipid Concentrate (Thermofisher Scientific, Cat. no. 11905031) and 1X Penicillin-Streptomycin (Thermofisher Scientific, Cat. no. 15140122)) and placed on orbital shaker. From day 18 on media change was performed every three-four days. On day 35, the media was changed to CDMIII (DMEM/F12 Glutamax supplement (Thermofisher Scientific, Cat. no. 31331093) with 10% FBS (GE Healthcare Life Sciences, Cat. no. SH30070.03), 5 µg/mL Heparin (Merck, Cat. no. H3149-25KU), 1X N2 Supplement, 1X CD Lipid Concentrate and 1% Matrigel (Corning, Cat. no. 356234)). Organoids were cultured in CDMIII until day 50.

Starting from day 21 until the collection day, dorsal forebrain organoids were treated with Valproic acid sodium salt (Sigma Aldrich, Cat. no. P4543) and Levetiracetam (Sigma Aldrich, Cat. no. L8668). The treatment was supplemented into the culturing media for treatment groups and refreshed on every media change. For day 35 experiments, 0.1, 0.5 and 1 mM VPA and LEV were used individually for treatment groups, and for day 50 experiments, 1 mM VPA were used for treatment group. Information on treatment conditions can be found in the figures and figure legends.

### Size measurements

Bright field images of organoids were taken every three-four days for organoids treated from day 21 to day 35 and every five days for organoids treated from day 21 to day 50 starting from day 21. Images were taken with EVOS cell imaging system (Thermo Fisher) with 4x or 10x objectives and were analyzed using a previously published Fiji (Schindelin et al., 2012) macro (Ivanov et al., 2014) with minor modifications. Then, size measurements data was arranged in Microsoft Excel and plotted using GraphPad Prism.

### Immunohistochemistry (IHC) in dorsal forebrain organoid slices

On day 35 or day 50, organoids were collected and fixed in 4% paraformaldehyde (PFA) (Morphisto, Cat. no. 11762) for one hour at room temperature. Fixed samples were washed three times with PBS (Roth, Cat. no. 1105.1) for 15 minutes before incubation at 4°C with 30% sucrose (Sigma Aldrich, Cat. no. S7903) in PBS. Once the organoids settled down in the plate, they were embedded in 1:1 (v/v) mixture of optimal cutting temperature OCT compound (Sakura, Cat. no. 4583) and 30% sucrose in PBS solution to be frozen at −80°C freezer. The organoid blocks were sectioned to 20 µm-thick slices on Superfrost Plus slides (VWR) with a cryostat and kept in −80°C freezer until further use.

For immunohistochemistry, the slides were thawed in room temperature for 15 minutes and rehydrated with PBS. After removal of extra embedding, hydrophobic pen (PAP pen, Abcam, Cat. no. ab2601) was used to circle the sections. Sections were incubated with permeabilization and blocking solution (1% Triton-X100 (Sigma, Cat. No. T8787), 0.2% gelatin (Sigma, Cat. no. G1890), and 10% normal donkey serum (Abcam, Cat. no. ab7475) in PBS) for one hour in room temperature. Primary antibodies were diluted in permeabilization and blocking solution and incubated overnight at 4°C. The next day, primary antibodies (anti-Ki-67 (rabbit, 1:300, Merck, Cat. no. AB9260), anti-SOX2 (goat, 1:500, R&D Systems, Cat. no. AF2018), anti-CTIP2 (rat, 1:400, Abcam, Cat. no. ab18465), anti-VIM (mouse, 1:250, Millipore, Cat. no. V6630), and anti-cCas3 (rabbit, CST, Cat. no. 9661S)) were washed three times with PBS for 15 minutes. Secondary antibodies (donkey anti-goat AF555 (Abcam, Cat. no. ab150130), donkey anti-goat AF647 (Invitrogen, Cat. no. A21447), donkey anti-rat AF555 (Abcam Cat. no. ab150154), donkey anti-rabbit AF488 (Invitrogen Cat. no. A21206), donkey anti-rabbit AF647 (Invitrogen, Cat. no. A31573), donkey anti-mouse AF488 (gifted)) were diluted in permeabilization and blocking solution (1:1000) and incubated for three hours at room temperature. Secondary antibodies were washed three times with PBS for 15 minutes and stained with DAPI (Thermofisher Scientific, Cat. no. D1306) diluted in PBS (1:5000) for four minutes at room temperature. After rinsing with PBS, the slides were mounted with ProLong Gold (Thermofisher Scientific, Cat. no. P36930). For epifluorescence imaging of whole organoid sections, Leica DMi8 microscope was used with 20x or 40x magnification objectives. For confocal imaging of VZ-like structures, Zeiss LSM710 confocal microscope, Leica SP8 confocal microscope was used with 40x magnification objective.

### Quantification of IHC analysis

Imaging of Ki67+ cells were performed on randomly selected VZ-like regions in the dorsal forebrain organoid slice. Using Imaris 9.7 software, all confocal images were rotated, adjusted with “Surfaces” tool and cropped to obtain 50 µm-wide VZ-like regions. Then, by using “Spots” tool cells that were positive for both Ki67 and SOX2 markers were quantified. The exact number of VZ-like regions quantified for each treatment condition can be found in the respective figure legends.

Quantification of the number and size of VZ-like structures were performed with ImageJ. The whole DFO section was used to count the total number and area of VZ-like regions. Polygon selection tool was used to mark each VZ-like region using SOX2+ vRG cells, then total number of polygons created, and the area of each polygon was quantified in ImageJ.

Quantification of cCas3+ area over SOX2+ VZ-like area was performed using custom macro in ImageJ. In short, first VZ-like regions were cropped in ImageJ using SOX2+ vRG cells as reference. The macro would first detect the SOX2+ area within the VZ-like region and crop the image where SOX2 signal is positive, then identify the cCas3+ and SOX2+ double positive area to generate percentage cCas3+ area over SOX2+ VZ-like region.

Quantification of Vimentin signal was performed using custom macro in ImageJ. In short, segmented line selection tool with defined width was used to mark the edge of all the organoid images taken at 20x magnification. Then, mean gray value in the selected region was quantified for vimentin signal at the organoid edges.

### RNA isolation and bulk RNA sequencing

RNA isolation from organoids was performed by using the RNeasy Mini Kit (Qiagen, Cat. No. 74106) and RNase-Free DNase set (Qiagen, Cat. no. 79254) with RNAase-free water. Library preparation for bulk RNA sequencing was performed by Novogene (https://en.novogene.com/) using non-directional poly-A enrichment strategy. Samples were sequenced with Illumina NovaSeq 6000 platform using paired-end, 150-bp-long reads to 30 million reads per sample. Further quality control measures and bioinformatics analysis were performed by Novogene.

For detection of significantly differentially expressed genes (DEGs) from differential gene expression analysis, two criteria were used: (1) absolute Fold change > 1.5 and (2) padj ≤ 0.05 between Ctrl and VPA treatment conditions. For further GO enrichment analysis of upregulated and downregulated DEGs, online RNA-seq bioinformatics analysis tool of Novogene called NovoMagic was used. All expressed genes present in the dataset was used as background for enrichment analysis. Further plotting of the data was performed on R using custom scripts.

Layer-specific analysis and plotting was performed using custom R script following the method published in Miller *et al. (32).* For enrichment human fetal PCW15-16 microarray data from Allen Institute was used. Fisher’s exact test was used for statistics with the Benjamini-Hochberg for multiple comparisons.

Weighted gene co-expression analysis (WGCNA) was performed by Novogene. NovoMagic was used for further GO enrichment analysis of eigengene modules generated by Novogene. Custom script in R were used to visualize GO analysis and WGCNA module network using the gene names as nodes and connectivity between genes as size of each node.

## Single cell RNA sequencing

### Single cell dissociation and multiplexing for single cell RNA sequencing

Following a published protocol (Velasco et al.) with minor modifications, individual dorsal forebrain organoids were dissociated into single cells using Worthington Papain Dissociation System kit (Worthington Biochemical, Cat. no. LK003150). In short, organoids of interest were washed twice with PBS and then incubated with Papain supplemented with DNase inhibitor for 20 mins at 37°C on orbital shaker. Then, organoids were gently triturated 15 times using 1 mL pipette tip and returned to incubation 10 minutes, repeated twice. Single cells were transferred to a tube containing Ovomucoid Inhibitor diluted in Earle’s medium supplemented with DNase inhibitor and centrifuged at 150 g for 10 minutes in room temperature. Cell pellet was resuspended in 0.04% BSA in PBS, filtered through 40 µm cell strainer (Flowmi) and counted using an automatic fluorescent cell counter (Luna). Single cell suspensions were centrifuged at 300 rcf for 5 minutes at room temperature. The cell pellet was resuspended in Cell Multiplexing Oligos from 3’ CellPlex Kit Set A (10x Genomics, Cat. no. PN-1000261) and multiplexing protocol from manufacturer was performed.

Finally, multiplexed cells were counted and pooled in equal proportions to be proceeded with single cell RNA sequencing pipeline by CeGat Tübingen. Approximately 14 000 cells per channel were loaded to Chromium chip (10x Genomics) and processed through the Chromium controller. Sequencing libraries were generated using Chromium Next GEM Single Cell 3’ Kit, v3.1 (10x Genomics) and sequenced on NovaSeq 6000 (Illumina).

## Single-cell data analysis

### Quality control and data preprocessing

CellRanger (v.7.1.0) multi pipeline was used to align reads to the prebuild 10x Genomics GRCh38-2020-A reference genome including introns and generate cell-by-gene matrices. Data were analyzed in R (v4.3.2) using Seurat (5.0.3)(33). Ambient RNA was removed with CellBender (v0.2.0) with false discovery rate (FDR) = 0.01 and epochs = 150 (34). The following thresholds were chosen for removing low-quality cells: min number of genes per cell – 2000, max number of reads per cell – 50000, max percentage of mitochondrial reads per cell – 8%, min percentage of ribosomal protein coding genes – 4%, min log10(genes per UMI) – 0.8. DoubletDetection (v4.2) under default parameters was employed to remove potential doublets (35). Following these filters, a total of 37,326 cells were retained with an average of 16,842 transcripts per cell and average 5,159 genes per cell. Raw gene counts were normalized per sample using the SCTransform workflow while regressing out the number of genes, the number of reads, and percentage of mitochondrial and ribosomal RNAs. Different samples were then integrated using reciprocal principal component analysis as implemented in Seurat v5. Linear dimensionality reduction was performed using PCA, and we selected first 30 PCs for further clustering and non-linear dimensionality reduction based on the Elbow plot. UMAP plots were generated using the Seurat package with a number of neighbors set to 15. Louvain clustering was performed across several resolutions, and resolution 0.6 was chosen based on clustree (0.5.1) cluster stability assessment (36). Cluster marker analysis using the FindAllMarkers function with both MAST (37) and roc algorithms were utilized as an initial guidance for cluster annotations. Additionally, we assessed the expression of canonical genes to assign each cluster to a known cell type.

### Removal of non-telencephalic and stressed cells

The VoxHunt algorithm was used to assess the regional identity of individual clusters, and SCTransform expression values were used as input. E18 mouse transcriptional profiles were used as a reference for identifying regional identity. A cluster that did not show similarity to the forebrain was filtered from the dataset.

After excluding non-telencephalic cells, the Gruffi (1.5.5) algorithm was applied to identify and remove cells exhibiting a transcriptional signature of cellular stress. This was done using an automatic threshold at the 0.9 quantile for two stress-related GO terms - GO:0034976 (Response to ER stress) and GO:0006096 (Glycolytic process) - and two non-stress-related GO terms - GO:0042063 (Gliogenesis) and GO:0022008 (Neurogenesis). After removing non-telencephalic and stressed cells, the dataset was subjected to normalization, integration, clustering (with resolution 0.4), dimensionality reduction, and annotation workflow as mentioned above.

### Cell proportion analysis

Differences in cell type proportions between experimental conditions were analyzed with a permutation test followed by bootstrapping (https://github.com/rpolicastro/scProportionTest), where clusters with absolute log2 Fold change > 0.58 and FDR < 0.05 were considered differentially abundant.

### Trajectory inference and trajectory quality assessment

For this section and subsequent steps of the analysis of the single-cell RNA-seq data, updated versions of R (v.4.4.1) and Seurat (v.5.1.0) were utilized. For trajectory inference, cells belonging to excitatory lineage were subset, normalized, integrated, and plotted into reduced dimensions. Monocle 3 (v.1.3.7) algorithm was applied to construct putative single-cell differentiation trajectories based on the UMAP representation (38, 39). The start of the trajectory was set in the RG cell type. A random forest classifier was used to retrieve genes differentially expressed along the putative deep-layer neuron differentiation trajectory.

### Identification of differentially expressed genes, gene module signatures, and GO enrichment analysis

To identify differentially expressed genes (DEGs) between experimental conditions by cell type, we performed a differential gene expression (DGE) analysis using the edgeR package (v.4.2.2). A minimum threshold of 30 cells per sample for each cell type was applied. Three pseudo-replicates were generated for each sample and cell type by splitting the cells into three groups and summing the gene expression values within each group. Low-expression genes were excluded using the filterByExpr function, and differences in library sizes across samples were normalized with calcNormFactors. Dispersion estimates for each gene were calculated using estimateDisp, and a negative binomial generalized log-linear model was fitted to the gene counts with glmQLFit. Genes with absolute log2 Fold change > 1.5 and FDR < 0.0001 and were considered statistically significant.

GO enrichment analysis was conducted for both upregulated and downregulated genes using the enrichGO function from the clusterProfiler R package (v.4.12.6), with an adjusted *p*-value cutoff of 0.05 and the false discovery rate (FDR) correction method. Genes expressed in the dataset were used as the background for enrichment calculations.

To evaluate differential expression of gene modules, the SCTransform-normalized count matrix was used as input for the UCell algorithm (v.2.8.0). Predefined gene sets corresponding to selected GO terms were employed to compute enrichment scores at the single-cell level.

Gene set enrichment analysis was also performed for autism spectrum disorder (ASD) risk genes obtained from the SFARI database (SFARI released on 31.10.2019, 20.07.2022 and 09.10.2024)(40). Enrichment was evaluated using a one-sided Fisher’s exact test in R, which assessed whether the proportion of ASD risk genes within the differentially expressed gene set for each cell cluster was significantly higher than expected by chance.

### Cell Communication Analysis

We employed CellChat (v.2.1.2) to analyze cell-cell and cell-ECM interactions across experimental conditions. Seurat objects were split by condition, and for each subset, normalized expression data and metadata were used to create CellChat objects. The CellChatDB human database was subset to include ECM-receptor and cell-cell interactions separately, and overexpressed genes and interactions were identified for each condition. Communication probabilities were computed using the tri-mean method and filtered to exclude interactions involving fewer than 10 cells. Finally, aggregated signaling networks were generated, and interaction networks were compared across conditions to rank signaling pathways and evaluate differences in total interaction strength.

### Gene Regulatory Network Analysis

SCENIC analysis was performed to identify gene regulatory networks in RG cells, comprising a subset of 6210 cells extracted from the Seurat object containing excitatory lineage cells. The regulatory network inference step was conducted using the PySCENIC <u>(v.0.12.1)</u> grn command. Co-expression modules were calculated based on transcription factors in the human genome (41).

The PySCENIC ctx command was subsequently employed to annotate and refine the regulatory network. Regulatory modules were filtered by motif enrichment using two motif databases: hg38 refseq-r80 10kb_up_and_down_tss.mc9nr and hg38 refseq-r80 500bp_up_and_100bp_down_tss.mc9nr (42). Motifs with a normalized enrichment score (NES) ≥ 3.0 and either direct transcription factor annotations or orthologous annotations were used to define regulons, with a minimum threshold of 10 genes per regulon. AUCell analysis was carried out using the PySCENIC aucell function to calculate the activity of each inferred regulon across individual RG cells. Additionally, regulon specificity scores were computed to evaluate the condition-specific activity of regulons within RG cells. For visualization, a t-SNE embedding was generated using the AUCell regulon activity matrix, and specific regulon activity levels were displayed using Scanpy’s plotting functions.

## Mass spectrometry sample collection and protein isolation

Whole organoid proteome and secretome samples were collected as organoids or their culturing media, respectively, from the same organoids. Within each sample, 3 organoids or 3 media aliquots from respective organoids were pooled. For secretome analysis, media change was performed on D19, D34, and D49 to collect one-day old media on D20, D35, and D50. Media were collected into a 2 mL tube, submerged in liquid nitrogen until frozen, and stored at −80°C. For D49 media change, CDMIII media without HyClone Defined Fetal Bovine Serum (Cytiva, Cat. no. SH30070.03) and Matrigel was used to avoid artifacts in protein composition. For whole organoid proteome analysis, after collection of the culturing media, dorsal forebrain organoids on D20, D35, and D50 of differentiation were washed with 1X PBS, then transferred to protein LoBind 1.5 mL tubes (Eppendorf, Cat. no. 0030108116). Once transferred, any liquid was removed, and tubes containing organoids were submerged in liquid nitrogen for snap freezing and stored at −80°C until protein isolation. The samples were delivered to the Core Facility for Medical Bioanalytics for protein isolation and mass spectrometry.

Total protein was extracted from collected samples using TBS lysis buffer (Tris-(hydroxymethyl)-aminomethane (30mM; Tris (AppliChem), NP40 0.5%, supplemented with complete protease inhibitors cocktail (Roche, Cat. no. 11836170001) and phosphatase inhibitors 2 and 3 cocktails (Sigma Cat. no. P5726 and Cat. no. P004) according to the manufacturer recommendations. Total extracted proteins were precipitated and quantified using Bradford assay, and a similar total amount of protein from each sample was used to prepare the sample for mass spectrometry analysis as described previously (43).

For secretome, an equal amount of secretome collected from different biological conditions were centrifuged at 5,000 g for 10 minutes at 4°C to remove cell debris and subjected to a methanol/chloroform precipitation as adapted from a published study (44). Briefly, the collected supernatant volume was mixed with four volumes ice-cold methanol, vortexed and centrifuged for one minute at 9,000 g. Then, mixed with one volume of chloroform, vortexed and centrifuged for one minute at 9,000 g. Three volumes of HPLC grade water were added, vortexed and centrifuged for two minutes at 16,000 g. After discarding the upper phase, without disturbing the interphase, three volumes of HPLC grade water were added, vortexed and centrifuged at 16,000 g for four minutes. Finally, the supernatant was discarded, and the protein pellets were dried under a laminar flow hood and stored at −80°C to be processed for in solution tryptic digestion.

Briefly, extracted proteins were alkylated and reduced using dithiothreitol (DTT) and iodoacetamide (IAA), followed by tryptic digestion overnight at 4°C. Collected peptides were further desalted using STAGE-tip (Affinisep). Peptides were resuspended in Acetonitrile trifluoracetic acid solution and prepared for Mass spectrometry analysis.

## Mass Spectrometry

Mass Spectrometry analysis was performed on an Ultimate3000 RSLC system coupled to an Orbitrap Tribrid Fusion mass spectrometer (Thermo Fisher Scientific). Tryptic peptides were loaded onto a µPAC Trapping Column with a pillar diameter of 5 µm, inter-pillar distance of 2.5 µm, pillar length/bed depth of 18 µm, external porosity of 9%, bed channel width of 2 mm and length of 10 mm; pillars are superficially porous with a porous shell thickness of 300 nm and pore sizes in the order of 100 to 200 Å at a flow rate of 10 µl per min in 0.1% trifluoroacetic acid in HPLC-grade water. Peptides were eluted and separated on the PharmaFluidics µPAC nano-LC column: 50 cm µPAC C18 with a pillar diameter of 5 µm, inter-pillar distance of 2.5 µm, pillar length/bed depth of 18 µm, external porosity of 59%, bed channel width of 315 µm and bed length of 50 cm; pillars are superficially porous with a porous shell thickness of 300 nm and pore sizes in the order of 100 to 200 Å by a linear gradient from 2% to 30 % of buffer B (80% acetonitrile and 0.08% formic acid in HPLC-grade water) in buffer A (2% acetonitrile and 0.1% formic acid in HPLC-grade water) at a flow rate of 300 nl per min. The remaining peptides were eluted by a short gradient from 30% to 95% buffer B; the total gradient run was 120 min. Spectra were acquired in Data Independent Acquisition (DIA) mode using 50 variable-width windows over the mass range 350-1500 m/z, MS2 scan range was set from 200 to 2000 m/z.

## Mass spectrometry data analysis and statistics

MS RAW data were analyzed using DIA-NN 1.8.1 (PMID: 31768060) in library-free mode against the human database (UniProt release September 2023). First, a precursor ion library was generated using FASTA digest for library-free search in combination with deep learning-based Spectra prediction. An experimental library generated from the DIA-NN search was used for cross-run normalization and Mass accuracy correction. Only high-accuracy spectra with a minimum precursor false discovery rate (FDR) of 0.01, and only tryptic peptides (2 missed Tryptic cleavages) were used for protein quantification. The match between runs option was activated and no shared spectra were used for protein identification.

Statistical analysis, including label-free quantification ratios (LFQ), and two-sided corrected permutation-based T-test (250 permutations and a minimum p-value of 0.05) to identify putative differentially abundant proteins between the two groups was done using the Perseus software suite version 1.6.15.0 (PMID: 27348712). The analytical platform Omics Playground (https://github.com/bigomics/omicsplayground) was used for proteomics data exploration and integration.

Principle component analysis (PCA) plots and volcano plots were generated using the built-in functions within the Perseus software. For PCA plot of proteome and secretome, data before data imputation was used. For the rest of the analysis, data imputation was performed separately for each cell line and time point by replacing missing values with values from normal distribution. Custom codes in RStudio software were generated for heatmaps and GO analysis. To analyze and visualize proteins, we used gene symbols. pheatmap library was used for heatmaps with “Euclidean” as clustering distance for rows and averaged log2 transformed LFQ values were used as input for the heatmaps. Finally, overrepresentation analysis (ORA) for GO terms was performed using topGO library and parameters used for enrichGO function were as follows: reference gene list=org.Hs.eg.db, minGSSize=10, maxGSSize=500, p value=0.05, q value=0.10 and p adjusted method = FDR. All proteins that were found in at least two samples were used as background for GO enrichment analysis. Further adjustments to the plots were made using Inkscape software.

## Statistical analysis

Statistical analyses were performed in GraphPad Prism software or by using custom scripts in R/Python. The details of each statistical analysis performed can be found in the respective figure legend.

## Results

### Dose-dependent alterations in dorsal forebrain organoids are evident upon exposure to VPA, but not Levetiracetam

With the aim of investigating how environmental factors affect neocortical development, we differentiated a range of healthy control iPSC lines into dorsal forebrain organoids (23). To investigate the effects of different AEDs and their concentration-dependent biological effects, we treated DFOs with VPA (teratogenic AED) and LEV (non-teratogenic AED) at concentrations of 0.1, 0.5, and 1 mM from D21 of differentiation until D35 (Fig. S1A). These concentrations are in the clinically relevant therapeutic range for epilepsy treatment (45), and similar ranges were used in other neural organoid studies (25-29). We chose to start treatment from D21 targeting neural progenitors and early neural differentiation, mirroring the increased risk for neurodevelopmental disorders upon exposure in the first trimester of pregnancy (46). Throughout the treatment period, organoid growth was monitored using brightfield microscopy (Fig. S1B). At D35, VPA-treated organoids were smaller than control (Ctrl) organoids upon 1 mM VPA treatment. However, this effect was not clearly observed at lower concentrations (Fig. S1C). No change in organoid size was observed at any LEV concentration at any time point (Fig. S1C). To determine whether the smaller size of VPA-treated organoids was due to reduced proliferation in the VZ-like regions, we assessed the proliferative state of ventricular radial glia (vRG) by quantifying Ki67-SOX2 co-expressing proliferative radial glia (RG) cells over SOX2+ total RG cells in VZ-like region cropped to 50 μm-width (Fig. S1D). We found fewer proliferative RG cells upon VPA treatment on D35 in a concentration-dependent manner, where 1 mM VPA-treated organoids had significantly fewer proliferative vRG cells (Fig. S1E). We did not observe any significant reduction in proliferative vRG cells at any LEV concentrations (Fig. S1E). To determine whether cell death also contributed to the smaller size of these organoids, we quantified the percentage of cCas3+ apoptotic vRG cell area over a 50 μm-wide SOX2+ VZ-like region area (Fig. S1F). No significant difference was observed between the VPA-treated and Ctrl groups, indicating that apoptosis of vRG cells does not drive the reduced size at D35 (Fig. S1G). Since RG cells are the main cell type driving proliferation and organoid growth at D35, we decided to focus solely on this cell type for proliferation and apoptosis analysis. Collectively, these results suggest that VPA treatment alters the neurodevelopmental trajectory of DFOs by reducing the proportion of proliferative cells, in line with other VPA-treated organoid and animal models (26, 28, 47). As expected, we did not see significant changes upon LEV treatment, which is known as a non-teratogenic AED treatment. To understand the teratogenic mechanisms of VPA treatment we solely focused on VPA-treated DFOs in further analyses.

### Long-term exposure to VPA disrupts DFO growth and alters the structure of ventricular-like zones

To investigate the long-term effects of VPA on DFOs, we decided to use 1 mM VPA, as this concentration successfully mimicked the developmental defects without inducing apoptosis in vRG cells. Hence, 1 mM VPA was continuously added to the organoid culture media from D21 to D50 to focus on the early neurogenesis (Fig. 1A). Organoid growth was monitored with bright-field microscopy throughout the treatment period (Fig. 1B) and revealed that Ctrl organoids exhibited continuous growth over the culture period, while VPA-treated organoids ceased growing around day 35 (Fig. 1C). Size comparison between VPA-treated and Ctrl organoids revealed significant differences at both D35 and D50 (Fig. 1D).

**Fig. 1:**
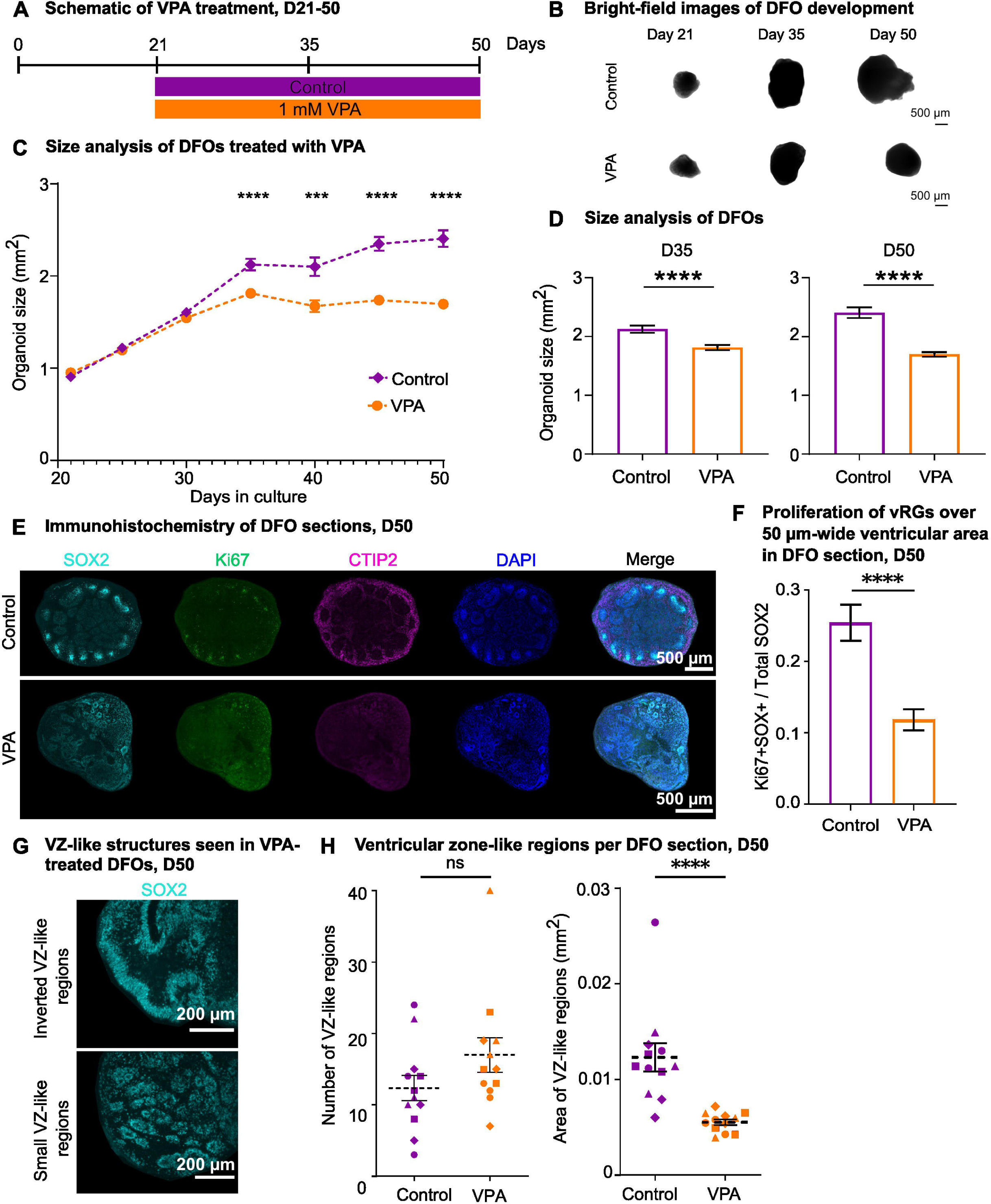
Dorsal forebrain organoids to study the effects of long-term exposure to VPA. **A.** Schematic of VPA treatment from day 21-50. **B.** Representative brightfield images of organoid development. **C.** Quantification of organoid size throughout treatment period with VPA compared to Ctrl. Data points represent the mean of all data in each condition, and the error bars represent the standard error of mean (SEM). For D21-35, n=72 for VPA and Ctrl; for D40 n=29 for VPA and n=30 for Ctrl; for D45-50 n=58 for VPA and n=59 for Ctrl. Multiple unpaired t-tests with Welch correction were used for significance testing between VPA and Ctrl per time point. Benjamini and Hochberg False Discovery Rate (FDR) was used for multiple comparisons. **D.** Bar plot of organoid size at D35 and D50. Bar plots represent the mean of all data points, and the error bars represent the standard error of mean (SEM). Welch’s t test was used for significance testing between VPA and Ctrl at D35. Nonparametric Mann-Whitney statistical test was used for significance testing at D50. Shapiro-Wilk test was used to assess normality. **E.** Immunohistochemistry of SOX2 for RG cells, Ki67 for proliferative cells and CTIP2 for deep layer neurons on representative whole organoid section. **F.** Quantification of proliferative RG cells in 50 µm-wide ventricular area in DFO slice at D50. Individual VZ-like areas from different organoids and batches are quantified, for Ctrl n=37 and for VPA n=45. Nonparametric Mann-Whitney statistical test was used for significance testing between VPA and Ctrl. Shapiro-Wilk test was used to assess normality. **G.** Representative images of disrupted VZ-like regions in VPA treated organoids. **H.** Quantification of number and area of SOX2+ VZ-like regions within DFO slice. For both number and area of VZ-like regions, n=12 organoids were used (n=3 per batch, 4 BIONi010-C batches). For area of VZ-like regions, all VZ-like regions per organoid were averaged. Nonparametric Mann-Whitney statistical test was performed for both number of VZ-like regions and area of VZ-like regions. Shapiro-Wilk test was used to assess normality. All data points were obtained from four independent DFO differentiation batches from BIONi010-C cell line. Statistical analysis, ****, p < 0.0001; ***, p < 0.001; **, p < 0.01; *, p < 0.05.

Furthermore, we visualized (Fig. 1E, S2A) and quantified the proliferative vRG using Ki67-SOX2 co-expressing cells over SOX2+ total RG cells at D50 (Fig. 1F). VPA-treated organoids had significantly less proliferating vRG cells compared to the Ctrl organoids at D50 (Fig. 1F).

A closer look at the SOX2-positive cells revealed that VZ-like structures were disrupted upon VPA treatment; for example, several DFOs displayed inverted VZ-like regions while others had small VZ-like regions (Fig. 1G). Overall, VPA-treated organoids showed a tendency to have more VZ-like structures, and the total area of VZ-like structures was significantly reduced at D50 (Fig. 1H). Due to the observed defects in the VZ-like structures, we performed intermediate filament cytoskeletal protein vimentin (VIM) antibody staining to evaluate the cellular architecture of the neural progenitor cells (Fig. S2B). This staining revealed an accumulation of VIM protein around the edge of the organoids upon VPA treatment compared to Ctrl, suggesting disrupted cytoarchitecture (Fig. S2C). Overall, our results corroborate previous findings that showed VPA-treated organoids had smaller VZ-like structures (28, 29).

### Transcriptomic analysis reveals that VPA predominantly affects the extracellular matrix, cell adhesion molecules, and synapses

To investigate the molecular changes driving the observed cellular features induced by long-term VPA treatment, we treated DFOs with 1 mM VPA from D21 to D50 and performed RNA sequencing (RNA-seq) on VPA-treated and Ctrl DFOs on D50 of differentiation. Principal component analysis (PCA) showed a clear separation of VPA-treated and Ctrl samples, indicating distinct expression profiles (Fig. 2A). Subsequently, we performed differential gene expression (DGE) analysis to compare gene expression between VPA treated and Ctrl organoids (significance threshold: absolute log2 Fold change > 0.58, p-adjusted value (padj) < 0.05) (Fig. 2B). DGE analysis identified 2,358 upregulated differentially expressed genes (DEGs) and 979 downregulated DEGs upon VPA treatment (Fig. 2B). Gene ontology (GO) enrichment analysis of all upregulated and downregulated DEGs showed upregulation of biological processes related to cell adhesion, secretion and extracellular matrix (ECM) organization and downregulation of biological processes (BP) related to neurogenesis and axogenesis upon VPA treatment (Fig. 2C). To investigate whether these DEGs corresponded to a specific cortical layer, we performed enrichment analysis of layer-specific genes by using human fetal postconceptional week (PCW) 15-16 microarray data (32) (Fig. 2D). We identified subplate and ventricular zone as cortical layers significantly susceptible to VPA treatment (Fig. 2D). BP related to proteoglycans in the subplate-specific DEGs and JAK-STAT signaling in the ventricular zone-related DEGs were downregulated (Fig. 2E). JAK-STAT signaling is necessary for cell cycle progression in some radial glia cells (48) and is shown to be affected also by other environmental impacts, such as pro-inflammatory cytokines, interleukin-6 (IL-6)(49).

**Fig. 2:**
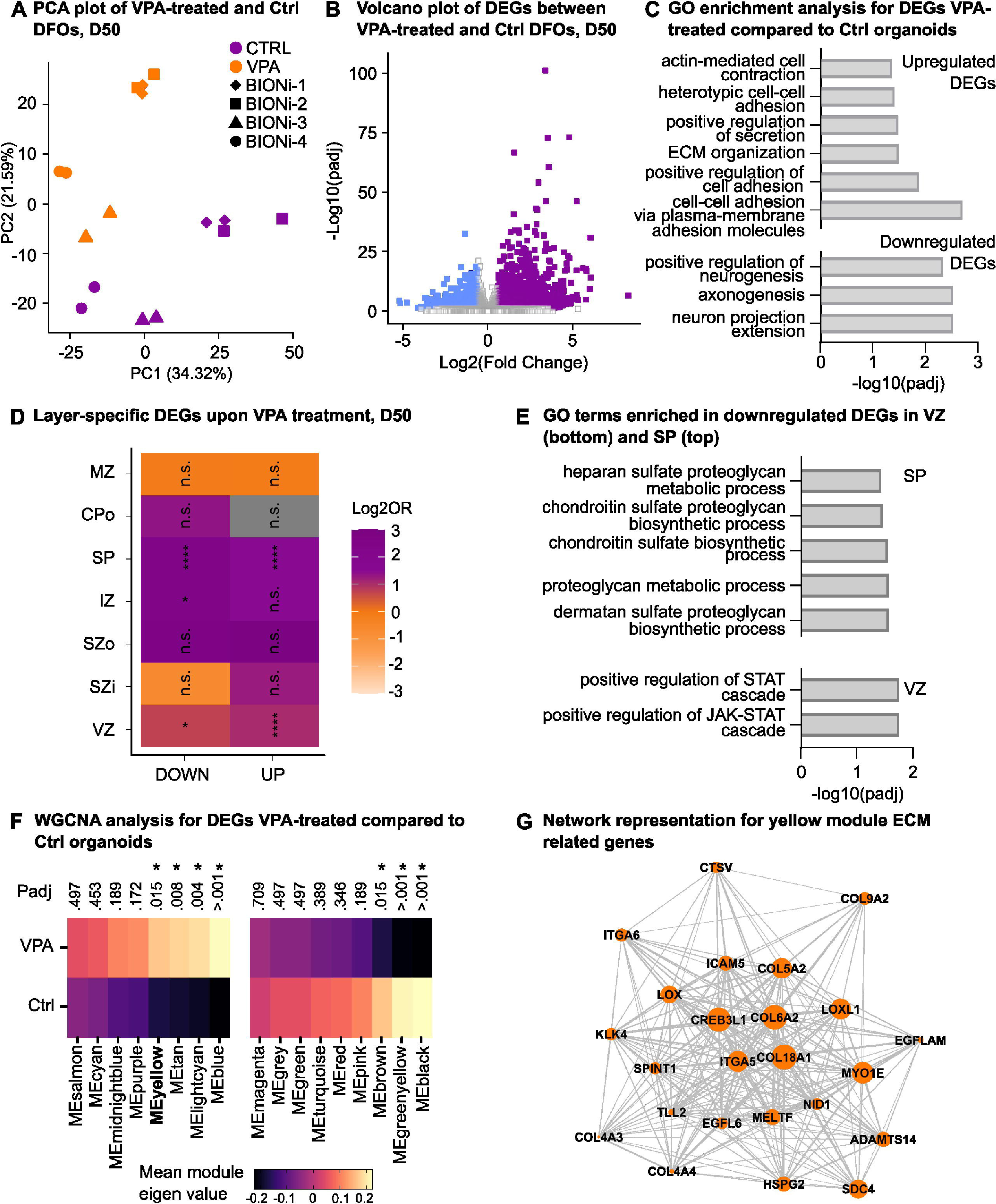
Bulk RNA-sequencing to identify changes in DFO transcriptome upon VPA exposure. **A.** PCA plot for DFO samples at D50. Color of the data point represents the treatment condition (purple, Ctrl; orange, VPA) and shape of the data point represents the organoid batch. Four batches generated with BIONi010-C cell lines were used for RNA-sequencing. **B.** Volcano plot of VPA vs. Ctrl. Significance threshold is absolute Fold change > 1.5 and padj from t-test < 0.05. Each datapoint represents a gene. Upregulated differentially expressed genes (DEGs) are colored in purple, downregulated DEGs in blue, and not DEGs in grey. **C.** Bar plot showing selected biological processes derived from GO analysis using both significantly upregulated and downregulated DEGs between VPA-treated and Ctrl DFOs. **D.** Enrichment of layer specific genes among DEGs upon VPA treatment were analyzed as described as Miller *et al.(32).* For enrichment human fetal PCW15-16 microarray data from Allen Institute was used. Fisher’s exact test is used for statistics with the Benjamini-Hochberg for multiple comparisons. SG, subpial granular zone; MZ, marginal zone; CPo, outer cortical plate; CPi, inner cortical plate; SP, subplate; IZ, intermediate zone; SZo, outer subventricular zone; SZi, inner subventricular zone; VZ, ventricular zone. **E.** Bar plot showing selected biological processes derived from GO analysis using VZ- and SP-enriched significantly downregulated DEGs between VPA and Ctrl DFOs. **F.** Expression profile heatmap from WGCNA analysis of VPA and Ctrl treatment groups obtained by average expression of module genes. After checking descriptive statistics and performing normality test, multiple unpaired t-test was performed for statistical analysis. **G.** WGCNA network representation for yellow module genes enriched in ECM-related GO terms.

Genes in biological systems do not function in isolation but in conjunction with other genes, and bioinformatic methods such as weighted gene co-expression network analysis (WGCNA) identify expression modules that group relevant genes using the correlated expression patterns (50). We hence performed WGCNA and identified 17 gene modules (Fig. S3A). From these modules, 7 were significantly deregulated upon VPA treatment (Fig. 2F). Among the significant modules, yellow, brown, and black modules supported the previous findings. GO analysis for the yellow module revealed an upregulation of cell-substrate adhesion and ECM organization-related terms (Fig. S3B), while the brown module showed a downregulation of axogenesis and synapse organization-related terms (Fig. S3C). The black module showed a downregulation of protein translation and endoplasmic reticulum (ER)-related terms (Fig. S3D). Downregulation of protein translation was also reported in DFO model of maternal immune activation upon IL-6 treatment (49). Network connection of selected ECM-related genes in the yellow module showed a high number of connections between collagen, integrin, and metallopeptidase genes and other genes in the module, suggesting that defects in these genes might have an impact on the rest of the module (Fig. 2G). Overall, these findings highlight significant disruptions in ECM organization and cell adhesion pathways in response to VPA treatment and indicate a possible susceptibility of RG cells to VPA-induced effects. Significant enrichment in extracellular structure organization was also reported in transcriptomics analysis of VPA-treated organoids in other studies (25, 26), indicating a high reproducibility across independent studies using different organoid differentiation protocols and dosing schemes.

### Single-cell transcriptomic analysis suggests that VPA-treated DFOs have altered cell type proportions

Our immunohistochemistry (Fig. 1) and RNA-seq (Fig. 2) results indicate potential cell-type-specific effects of VPA treatment, especially targeting RG cells in VZ-like regions and newborn neurons in the subventricular zone in DFOs. To investigate cell-type-specific effects of VPA treatment, we performed single-cell RNA-seq (scRNA-seq) at D50 of differentiation on Ctrl and 1 mM VPA-treated DFOs (Fig. 3A). Each individual organoid was used as a sample for the analysis. The transcriptomes of 37,326 cells were analyzed after quality control, with on average 5,159 unique transcripts per cell (Fig. S4A). After performing normalization and sample-wise integration, the main cell types were annotated (Fig. S4B), and cells labeled as mixed inhibitory/excitatory neurons (mixed Inh/ExN), and non-forebrain cells were removed. Then, stressed cells were assessed using the Gruffi algorithm (51), by examining stress-, gliogenesis-, and neurogenesis-related GO terms (Fig. S4C). In total, we removed 10,827 cells, of which 1,719 cells were identified as mixed Inh/ExN or non-forebrain cells, and 9,108 cells were identified as stressed. The proportion of stressed cells between Ctrl and VPA-treated organoids was not pronouncedly different for any cell types (Fig. S4C). After removal of these cells, 14 transcriptionally distinct clusters were formed (Fig. S4D). UMAP plots showed that samples and cell lines were integrated (Fig. S4D). Their gene counts and ribosomal and mitochondrial gene counts were also visualized as quality control measures (Fig. S4E). We further annotated these clusters using manually selected marker genes (Fig. S5A) and confirmed the final annotations using reference-query integration with a human neocortical developmental dataset (52) (Fig. S5B). UMAP plot split by treatment conditions showed that VPA-treated organoids contained more progenitor and early neuronal cells, while Ctrl organoids contained more mature neurons (Fig. 3B). We then quantified the proportion of each cell type in each condition and sample (Fig. 3C, Fig. S5C). Progenitor cell clusters intermediate progenitors (IPC) and RG were overrepresented in VPA-treated organoids, and mature deep excitatory neurons (dExN) and inhibitory neurons (InhN) were overrepresented in Ctrl organoids (Fig. 3D). As a next step, we performed Monocle3 trajectory analysis using the excitatory neuron lineage (Fig. 3E). We observed the proportion of cells from VPA-treated organoids to be higher at earlier pseudotime, while the proportion of cells from Ctrl organoids was higher at later pseudotime (Fig. 3F). Cell type proportions and trajectory inference analysis showing reduced numbers of more differentiated neuronal cell populations was in line with downregulation of neurogenesis and synaptogenesis-related terms observed in the RNA-seq dataset. Altogether, these results suggest that VPA-treated organoids were stalled in their development compared to Ctrl organoids.

**Fig. 3:**
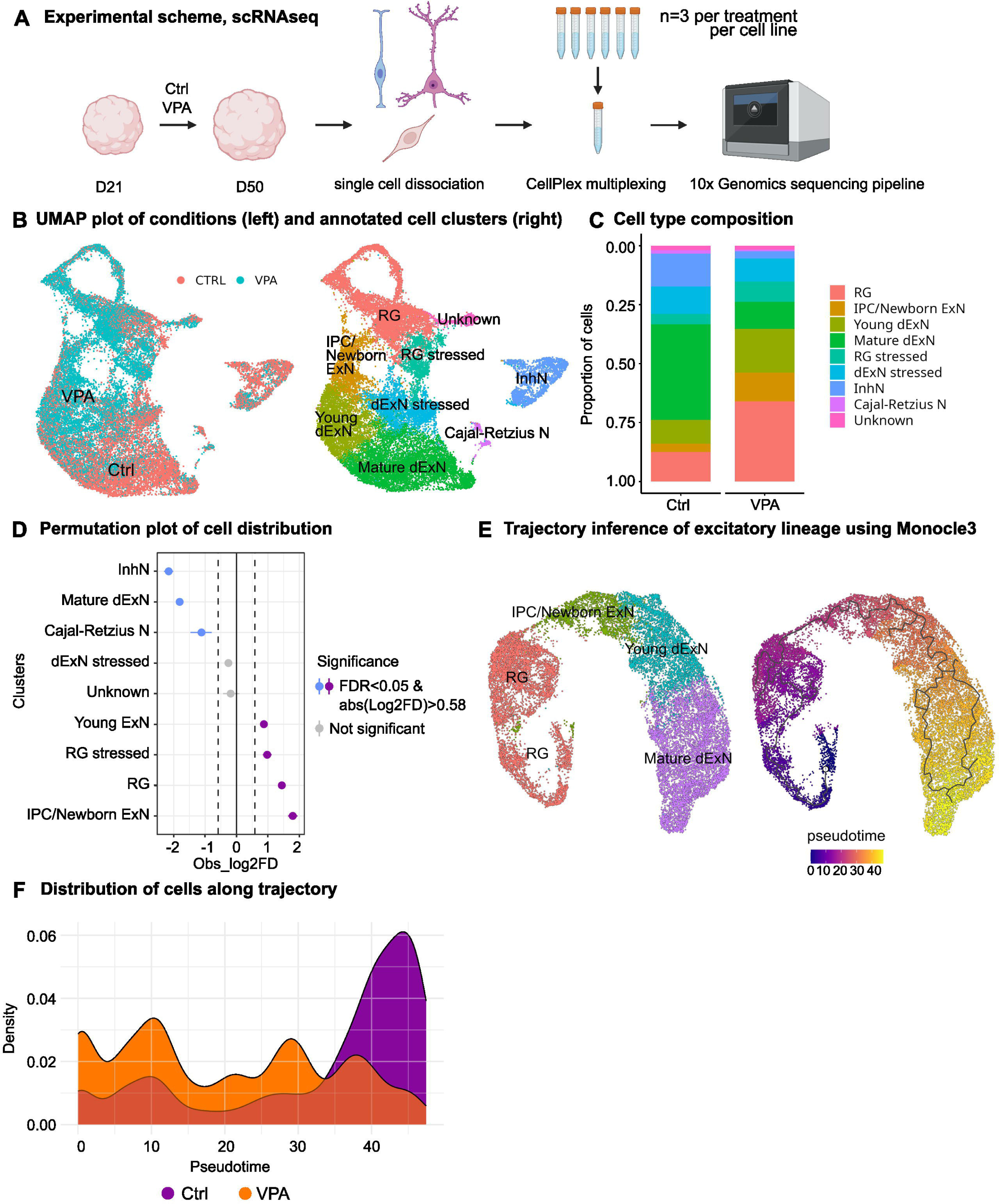
Single cell RNA-sequencing to determine affected cell types upon VPA treatment. **A.** Experimental scheme for VPA treatment of DFOs and single cell RNA sequencing. **B.** UMAP plot with cells from VPA (cyan) and Ctrl (red) DFOs (left). UMAP plot with 9 cell groups labeled by color (right). Data from n=3 organoids per condition per cell line, 2 cells lines were used (KOLF2.1J and BIONi010-C). RG, radial glia; IPC/Newborn ExN, intermediate progenitor cell/newborn excitatory neuron; Young dExN, young deep-layer excitatory neuron; Mature dExN, mature deep-layer excitatory neuron; InhN, inhibitory neuron; Cajal-Retzius N, Cajal-Retzius neuron; RG stressed, stressed radial glia; dExN stressed, stressed deep-layer excitatory neuron; Unknown. **C.** Stacked bar plot of proportion of cell types per condition. Cell lines were combined, n=6 per condition. **D.** Permutation plot for cell type distribution. Negative values represent underrepresented cell types in VPA-treated DFOs, while positive values represent overrepresented cell types in VPA compared to Ctrl DFOs. Significantly overrespresented cell types are colored in purple and underrepresented cell types in blue (significance threshold: absolute log2 Fold change > 0.58 and FDR < 0.05) and nonsignificant cell types are colored in grey. **E.** Trajectory inference of cells belonging to the excitatory lineage (including RG, IPC/Newborn ExN, Young dExN, Mature dExN) as assessed with Monocle3. The trajectory was started at the cycling cells, which corresponds to lower pseudotime and finished at Mature dExN cell cluster. **F.** Density plot of cell distribution along the pseudotime as assessed with Monocle3 and by experimental conditions, VPA and Ctrl.

### Different cell types show differential susceptibility to VPA treatment

To identify cell-type-specific changes in the transcriptome upon VPA treatment, we performed DGE analysis on transcriptomic clusters. We excluded Cajal-Retzius cells because they did not meet the threshold of a minimum number of 30 cells per organoid and performed DGE analysis on the remaining cell types (Fig. 4A). We identified more than 400 upregulated DEGs in RG cells, IPC/newborn ExN, young dExN, and mature dExN (significance threshold: absolute log2 Fold change > 1.5 and false discovery rate (FDR) < 0.0001) (Fig. 4B, 4C). We then performed enrichment analysis using upregulated and downregulated DEGs for each cell type. The young ExN cluster had significant upregulation of GO terms related to Wnt signaling pathway. Meanwhile, the mature dExN cluster had significant upregulation of GO terms related to dorsal/ventral patterning. Upon VPA treatment, neither yielded significant downregulated biological processes GO terms. For further analysis, we decided to focus on the progenitor cell types, RG and IPC, as these cell types were significantly over-represented in VPA-treated organoids. 426 genes were upregulated, and 115 genes were downregulated in RG in organoids treated with VPA compared to the Ctrl organoids (Fig. 4D). Overrepresentation analysis (ORA) using upregulated DEGs from RG cells identified upregulation of BP terms related to cell adhesion, synaptic membrane adhesion, and exocytosis-related terms in VPA-treated organoids compared to Ctrl (Fig. 4E). On the other hand, 416 genes were upregulated, and 109 genes were downregulated in the IPC/newborn ExN cluster in VPA-treated organoids compared to the Ctrl organoids (Fig. S5D). Similar to the RG cluster, ORA using upregulated DEGs from IPC/newborn ExNs identified BPs related to cell adhesion, synapse organization, vesicle fusion, and exocytosis in VPA-treated organoids compared to Ctrl (Fig. S5E). Downregulated DEGs from both RG and IPC/newborn ExNs did not yield any significant GO terms.

**Fig. 4:**
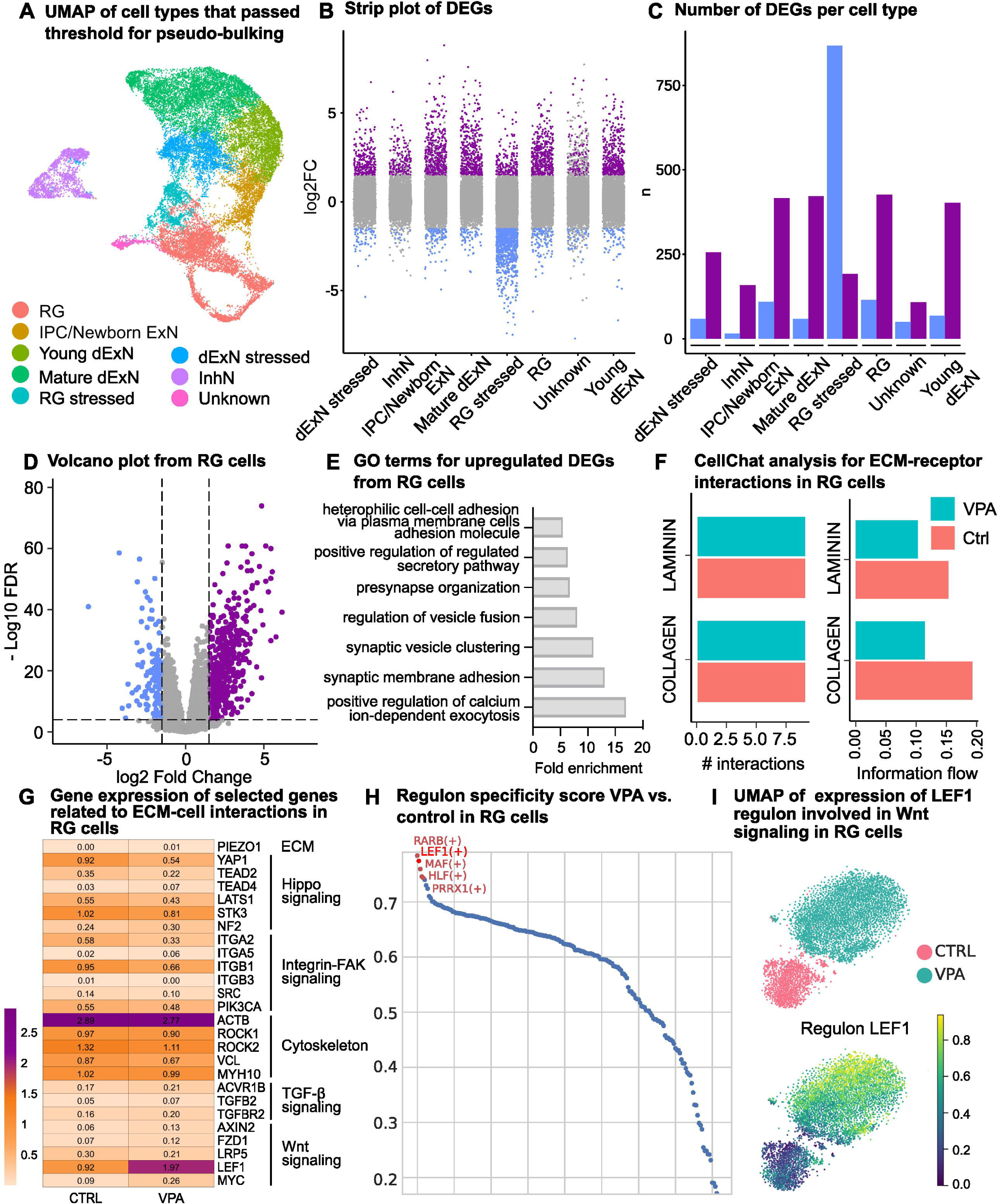
Single cell RNA-sequencing to analyze cell-type differential effects of VPA treatment. **A.** UMAP plot of cell types included in the DGE analysis. Cajal-Retzius neurons were not included due to not meeting the minimum cell number threshold for the analysis. RG, radial glia; IPC/Newborn ExN, intermediate progenitor cell/newborn excitatory neuron; Young dExN, young deep-layer excitatory neuron; Mature dExN, mature deep-layer excitatory neuron; InhN, inhibitory neuron; RG stressed, stressed radial glia; dExN stressed, stressed deep-layer excitatory neuron; Unknown. **B.** Strip plot of differentially expressed genes across each cell type (significance threshold: absolute log2 Fold change > 1.5 and FDR < 0.0001). Each datapoint represents a gene; upregulated DEGs are colored in purple, downregulated DEGs are in blue, nonsignificant genes are in grey. **C.** Bar plot of number of DEGs across cell types. Upregulated DEGs are colored in purple, downregulated DEGs are in blue. **D.** Volcano plot of DEGs in RG cells comparing the transcriptome of VPA-treated DFOs to Ctrl (significance threshold: absolute log2 Fold change > 1.5 and FDR < 1e-04). Each datapoint represents a gene; significantly upregulated DEGs are colored in purple, significantly downregulated DEGs are in blue, nonsignificant genes are in grey. **E.** Bar plot showing selected biological processes derived from GO analysis using significantly upregulated DEGs between RG cells of VPA-treated and Ctrl DFOs. **F.** CellChat analysis of ECM-receptor interactions comparing VPA (cyan) and Ctrl (red) transcriptomes of RG cells. Bar plot of number of interactions (left) and information flow (right). **G.** Heatmap of gene expression of selected genes from relevant signaling pathways; ECM, Hippo signaling, integrin-FAK signaling and cytoskeleton. **H.** Ranked regulons in RG cells and their specificity scores in VPA vs. Ctrl organoids as assessed by SCENIC. **I.** t-SNE plots obtained with SCENIC based on the AUCell regulon activity matrix in RG cells colored by condition (top) and showing the expression of the LEF1 regulon (bottom).

These results indicate altered extracellular secretion and intercellular communication in VPA-treated organoids. To explore this hypothesis in more detail, we used CellChat to compare two different types of molecular interactions in VPA vs. Ctrl: ECM-receptor interactions and cell-cell contact interactions in RG cells (Fig. 4F, Fig. S5F). In both cases, interaction strength was notably reduced in VPA compared to Ctrl organoids (Fig. 4F, Fig. S5F). We identified reduced information flow in ECM proteins such as collagen and laminin upon VPA treatment (Fig. 4F) and reduced information flow in synaptic adhesion molecules such as neurexin and NCAM (Fig. S5F). To understand the mechanism behind this change, we compared the expression of individual genes in the RG cluster from pathways that are likely to be involved in ECM interactions and cell adhesion (Fig. 4G). These pathways included PIEZO, Hippo signaling, integrin-FAK signaling, and cytoskeleton. We identified that upon VPA treatment, PIEZO1 channel expression was increased. Meanwhile, the overall expression of Hippo and integrin-FAK signaling pathway elements was decreased, suggesting changes in the ECM-receptor interactions in the RG cluster (Fig. 4G).

To further understand the regulatory mechanisms driving these signaling pathway changes, we performed SCENIC analysis on RG cells to identify key transcription factors driving transcriptional changes upon VPA treatment. We identified the 50 most active regulons in VPA-treated and Ctrl DFOs (Fig. S6A) and found RARB, LEF1, MAF, HLF, and PRRX1 to be highly specific to RG cells from VPA-treated organoids (Fig. 4H). Among these regulons, a canonical Wnt signal mediator LEF1 (53) had the second highest regulon specificity score and was highly expressed in RG cells from VPA-treated organoids (Fig. 4I). However, LEF1 was not significantly expressed in the RNA-sequencing dataset, which might indicate cell-type-specific changes in LEF1 expression.

Finally, to understand the relationship between VPA treatment and ASD, we tested for enrichment of cell-type-specific DEGs upon VPA treatment with ASD risk genes from SFARI (Fig. S6B). We identified the highest odds ratio for RG and IPC/Newborn ExN clusters, indicating that these cell types may be the most relevant for the increased ASD risk upon VPA exposure during pregnancy. This supports the idea that neural progenitor cells are the most susceptible cell types in DFOs to VPA treatment.

### Secretome analysis captures changes in ECM, CAM and synapses upon VPA treatment

Taken together, both bulk and single-cell transcriptomic data (Fig. 2-4, Fig. S3-6) point towards a disruption in the microenvironment, especially of progenitor cells, leading to the disorganized VZ-like structures we observed in immunohistochemical analysis (Fig. S2). To investigate the cellular microenvironment of VPA-treated and Ctrl DFOs, we performed secretome and proteome analysis on samples that consisted of three pooled organoid culture media or organoids in triplicates at D20, D35, and D50 using liquid chromatography-mass spectrometry (LC-MS) (Fig. 5A). Proteome comparison of VPA-treated and Ctrl DFOs yielded significant yet minor differences (Fig. S7A-D). For example, exocytosis and neuronal adhesion-related proteins such as α-synuclein (SNCA), annexin A6 (ANXA6) and astrotactin 1 (ASTN1) were increased in abundance upon VPA treatment at D35. In comparison, secretome comparison provided major significant findings. PCA analysis of all secretome samples showed three distinct clusters: D20 Ctrl DFOs, D35 and D50 Ctrl DFOs, and lastly D35 and D50 VPA-treated DFOs (Fig. 5B), suggesting a distinct secretome profile between VPA treatment and Ctrl conditions. To identify changes in protein abundance between conditions, we performed differential protein abundance analysis (significance threshold: absolute log2 Fold change ≥ 1, FDR < 0.05). The analysis of VPA-treated vs. Ctrl secretomes at D35 yielded 299 proteins with increased abundance and 387 proteins with decreased abundance (Fig. 5C, 5E). Meanwhile, the analysis of VPA-treated vs. Ctrl secretomes at D50 yielded 325 proteins with increased abundance and 547 proteins with decreased abundance (Fig. 5D, 5E). Functional annotation of these significantly differentially abundant proteins indicated a decrease in the BP GO terms related to synapse assembly and intermediate filament organization at D35 (Fig. 5F), and an increase in the BP GO terms related to cell adhesion, ECM organization, and collagen organization at D50 (Fig. 5G). A closer look at the selected ECM-related proteins and cell adhesion-related proteins showed an overall increase in protein abundance in both categories at D50 (Fig. 5I, 5J). To visualize the change in protein abundance over time, we plotted the protein abundance of COL5A1, COL9A1, and LAMA1 proteins at three time points for both VPA-treated and Ctrl secretomes (Fig. 5J). Time course plots showed an increased accumulation of ECM proteins over time in VPA-treated DFO secretomes compared to the Ctrl, which supports the findings from transcriptomic analysis that VPA treatment increases the ECM in the organoids (Fig. 2D). Taken together, our data indicates that altered VZ-like structures may result from disrupted interactions between the cellular microenvironment, cell adhesion molecules, and ECM proteins, which despite being abundantly secreted, do not appear to form functional interactions.

**Fig. 5:**
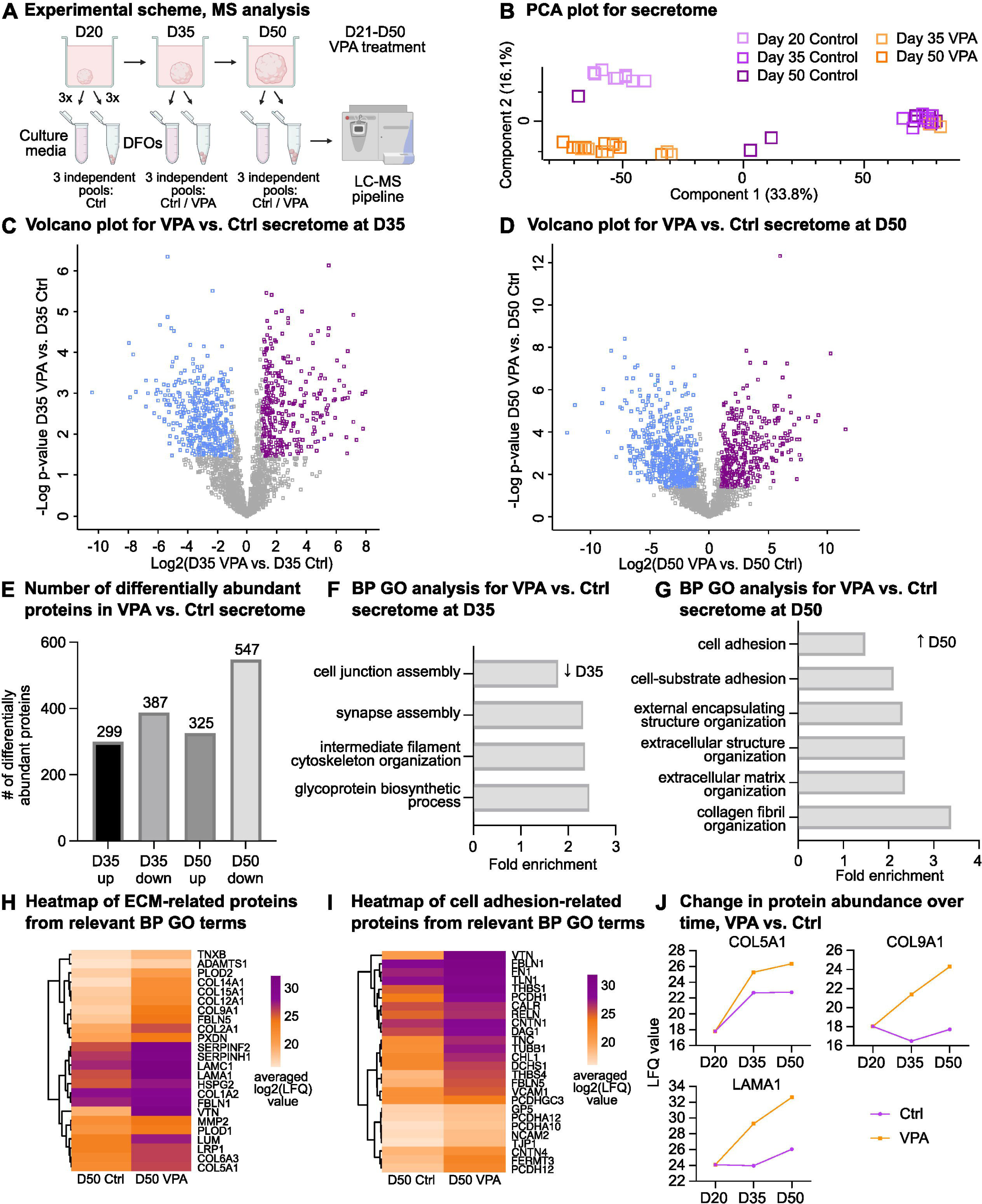
Secretome analysis of DFOs treated with VPA compared to Ctrl. **A.** Experimental scheme for collection of DFOs and DFO culture media for the proteome and secretome analysis, respectively. Three DFOs and three culture media were bulked to generate one pool, three independent pools were analyzed using liquid chromatography-mass spectrometry (LC-MS) at D20, D35 and D50. Same treatment scheme was used where DFOs were treated with 1 mM VPA from D21 until D50. BIONi010-C, KOLF2.1J and HMGU1 cell lines were used. **B.** PCA plot of all VPA-treated (orange) and Ctrl (purple) secretome samples from D20, D35 and D50. Each datapoint represents one sample, n=9 per time point. **C.** Volcano plot of differentially abundant proteins with increased abundance (purple) and decreased abundance (blue) proteins comparing VPA vs. Ctrl DFO culture media samples at D35. **D.** Volcano plot of differentially abundant proteins with increased abundance (purple) and decreased abundance (blue) proteins comparing VPA vs. Ctrl DFO culture media samples at D50. **E.** Bar plot showing number of differentially abundant proteins for VPA vs. Ctrl DFO culture media at D35 and D50. **F.** GO biological processes analysis of proteins with decreased abundance found in VPA vs. Ctrl secretome analysis at D35. **G.** GO biological processes analysis of proteins with increased abundance found in VPA vs. Ctrl secretome analysis at D50. **H.** Heatmap of selected ECM-related proteins from **G**. Colors represent the averaged log2 transformed LFQ values for VPA and Ctrl samples at D50. **I.** Heatmap of selected cell adhesion-related proteins from **G**. Colors represent the averaged log2 transformed LFQ values for VPA and Ctrl samples at D50. **J.** Line plot showing protein abundance over time for VPA-treated (orange) and Ctrl (purple) samples. Each point represents the averaged log2 transformed LFQ values at a given time for each condition. For **C**, **D** and **E** the difference between time points is plotted against -log(P-value) with significance threshold: absolute log2 Fold change values ≥ 1, FDR < 0.05).

## Discussion

Prenatal exposure to VPA is a well-known risk factor for neurodevelopmental disorders. This study mimicked early neurodevelopmental defects associated with prenatal VPA exposure using human iPSC-derived DFOs. Leveraging a multi-omics approach by combining several types of transcriptomic and proteomic analyses, we found an accumulation of ECM components in the microenvironment and dysregulation of mechanosensitive signaling pathways, especially in RGs. Hence, we highlight the link between intracellular dysregulation induced by VPA, most likely at the epigenetic level, and extracellular consequences and how they lead to the neurodevelopmental defects, an aspect that has been underexplored in previous human models of prenatal VPA exposure.

Previous studies on VPA-treated organoids have reported similar findings, including reduced organoid size and proliferation, and alterations in ventricular-like structures (26, 28, 29). A closer look at their transcriptomics data also identified upregulation of extracellular structure-related terms (25, 54). Altogether, despite differences in experimental paradigms, such as different organoid protocols, VPA concentrations, and treatment durations, these studies consistently reported similar neurodevelopmental disruptions, highlighting the robustness of VPA-induced defects. Similar to our findings, Zang et al. observed that VPA inhibited progenitor proliferation and reduced neural differentiation, as well as identified upregulation of genes related to the Wnt signaling pathway (26), which was also upregulated in our scRNA-seq data. They showed that blockers of Wnt pathway rescued VPA-induced defects such as surface area, ventricular-like structure thickness, neural progenitor proliferation, and neurogenesis (26). Interestingly, it was previously shown that the Wnt/β-catenin pathway is responsive to ECM stiffness (55, 56), which could link upregulated ECM genes to the increased expression of Wnt signaling genes observed by both Zang et al. and us. We therefore emphasize the role of the microenvironment in neurodevelopment and highlight how defects in extracellular interactions contribute to neurodevelopmental disorders.

### ECM accumulation in the cellular microenvironment and disrupted cellular mechanosensing

ECM is one of the main components of the cellular microenvironment and it provides mechanical and structural support to the cells as well as regulating the local availability of signaling molecules (57). ECM accumulation in the microenvironment may lead to increased ECM stiffness and thus influence cell-ECM interactions and cellular mechanosensing. Cell-ECM interactions can affect mechanosensing in two parallel pathways: integrin receptors and the integrin-based focal adhesion kinase (FAK) signaling pathway, and piezo channels (58, 59). Integrins are cell adhesion transmembrane proteins that link the ECM to the cytoskeleton, which allows them to transduce signals between cells and their microenvironment (60). This signaling is mediated by FAK, which is a key downstream signaling mediator of integrins upon activation (60). Integrin-FAK signaling components, including ITGA2, ITGB1, and ITGB3 as well as SRC were downregulated in VPA-treated DFOs, indicating a defective mechanotransduction (Fig. 4G). In line with our findings, it was previously shown that VPA treatment blocked the interaction between renal carcinoma cells and ECM by altering integrin expression and blocking FAK signaling (61, 62). VPA is a histone deacetylase inhibitor (HDACi), which alters expression of genes through epigenetic regulation (63). A possible epigenetic alteration of the integrin signaling pathway components upon VPA treatment can explain the decreased expression of integrin receptors, which may amplify ECM accumulation.

PIEZO1 expression was increased in VPA-treated DFOs (Fig. 4G), likely as a response to increased ECM in the microenvironment. Piezo channels are calcium channels that allow calcium flow into the cell depending on mechanical changes in the microenvironment (54). This calcium influx then activates various intracellular signaling pathways that influence processes such as growth and migration (64). In a preprint, ECM stiffening was shown to reduce proliferation, impair differentiation and induce accumulation of transitional alveolar progenitor cells via mechanosensing (65). These transitional cell states were associated with fibrotic remodeling and senescence (65). Thus, increased ECM components alter mechanical properties of the microenvironment and affect cellular mechanosensing.

### Hippo-YAP/TAZ signaling pathway dysregulation and quiescent/senescent cellular state

Different cellular inputs including mechanical signals, such as cell-cell contact and ECM stiffness can influence Hippo-YAP/TAZ signaling pathway, which regulates processes such as organ size by controlling proliferation/differentiation dynamics (66). In our dataset, disruptions in Hippo pathway components were observed, with reduced YAP1 and TEAD2 expression (Fig. 4G), which might indicate mechanosensing defects. It has been shown that YAP/TAZ are required to maintain the proliferative potential and structural organization of cortical RG cells (67). Meanwhile, loss of YAP/TAZ resulted in increased cell cycle length of RG cells, therefore reducing the proliferative capacity, and resulting in reduced number of cortical projection neurons (67). In line with this finding, reduced YAP1 expression in VPA-treated DFOs may explain overrepresentation of progenitor cells and underrepresentation of neuronal cells through increased cell cycle length or a possible shift towards transitional progenitor state due to incomplete RG to neuron differentiation.

Upon performing SCENIC analysis on RG cells, we identified upregulation of LEF1, a transcription factor associated with quiescent adult neural stem cell gene regulation (68). VPA could potentially increase the activity of LEF1 regulon by increasing its chromatin accessibility and drive radial glia cells into quiescence. Both quiescence and senescence are states of cell cycle arrest, and they have been proposed to represent a continuum of cell cycle withdrawal (69). Previously, decreased proliferation and increased senescent markers were observed in neuroepithelial cells in VPA-treated neural organoids (28) and VPA-treated neural tube organoids (70). ECM stiffening, such as through increased collagen, was shown to be one of the hallmarks of senescent cells (71). In line with this notion, our pseudotime analysis revealed an arrest at various cell states along the excitatory lineage (Fig. 3F). Thus, VPA might induce or prolong a transitional cellular state in progenitors, in which these cells then enter senescence or quiescence.

### Structural changes in VZ-like regions in DFOs and its neurodevelopmental implications

Altogether, our data reveals accumulation of ECM proteins in the microenvironment and defects in mechanosensing pathways, which likely result in disrupted VZ-like structures and associated neurodevelopmental implications. Consistent with our findings, Zhang et al. demonstrated that VPA exposure induces irregularly shaped neural rosettes with reduced diameters compared to Ctrl organoids modelling neural tube defects (72). Similarly, VPA treatment was used to model the risk of neural tube defects in human spinal cord organoids (73). Furthermore, VPA treatment of DFOs may be used to recapitulate the cortical malformations seen in ASD patients. For example, the reduced size of our DFOs upon VPA treatment could mimic microcephaly observed in some ASD patients (74). Therefore, *in vitro* VPA exposure of neural organoids offers an environmentally induced human model of ASD, providing a platform to study ASD pathophysiology in human tissue. Altogether, ECM components may be a therapeutic target for ASD, as modulation of the microenvironment has potential to restore molecular, cellular, and structural alterations. ECM-based interventions may offer a more accessible treatment option compared to treatments that target intracellular pathways.

In this study, we provide a comprehensive transcriptomic and proteomic profile of VPA-treated DFOs during early neural development. To our knowledge, this is the first neural organoid study to perform a comparative secretome analysis of VPA treatment in organoids. Our findings demonstrate that VPA treatment disrupts the microenvironment and cellular mechanosensing, potentially exacerbating their defects in a feedback loop. This disruption in ECM and cell-ECM interactions leads to impaired RG proliferation, defects in VZ-like structures, and disrupted neurogenesis. Therefore, this model can serve as a platform to study the effects of environmental adversities on the microenvironment and give new insights into the mechanisms of neurodevelopmental disorders.

### Limitations of the study

While we intentionally treated the organoids from D21 to D50 to capture changes in progenitor populations and early neuronal differentiation, this time frame does not provide information on later neurodevelopmental processes, such as neuronal maturation and synaptic activity. Additionally, although we employed a broad range of techniques, such as scRNA-seq and LC-MS, these methods have inherent limitations. For instance, scRNA-seq offers detailed insights into cellular states but lacks spatial context, while mass spectrometry is a bulk method that does not provide cell-type-specific proteomic changes or spatial information. In future studies, a new method called secretion encoded single-cell sequencing (SEC-seq) could be utilized to link the transcriptome of each cell with its secretome on a single-cell level (75). Furthermore, functional validation experiments are required to understand a possible causal relationship between ECM accumulation and changes in mechanosensing pathways following VPA treatment.

## Data availability

RNA sequencing, single cell RNA sequencing and MS data will be accessible in datatype-specific repositories.

## Acknowledgments

This study was funded by the WIN Program of the Heidelberg Academy of Sciences and Humanities, financed by the Ministry of Sciences, Research, and the Arts of the State of Baden-Württemberg (to SM) and Hertie Foundation (to SM). This project was also partially funded by the Brain & Behavior Research Foundation (NARSAD Young Investigator Grant 27026 to SM), the Baden-Württemberg state postgraduate fellowship (to KS and TK), Add-on Fellowship of the Joachim Herz Foundation (to KS), and the Daimler and Benz Foundation (32-06/20, to SM). We thank the German Research Foundation (DFG) for supporting the acquisition of the confocal microscope used in this study (INST 37/1170-1 FUGG, project number 467868227). This research has been partially funded by the Deutsche Forschungsgemeinschaft (DFG, German Research Foundation) under Germany’s Excellence Strategy via the Excellence Cluster 3D Matter Made to Order (EXC-2082/1 – 390761711 to SM and LB).

We thank Elisabeth Gustafsson for her technical support. We thank all the student assistants for their technical support. We thank Core Facility for Medical Proteomics, especially Franziska Klose, for protein isolation and mass spectrometry support. We thank Dr. Nicolas Snaidero and his lab members for the confocal microscopy support. We thank Dr. Christian Mahringer, Dr. Henner Koch and Prof. Dr. Yvonne Weber for fruitful discussions.

## Author contributions

ZY designed the study, performed experiments, data analysis, statistical analysis, and prepared the manuscript; KS, MAJ, TK, and KB performed experiments; KS, LB, CK, and MAJ performed data analysis; all authors revised the manuscript; SM conceived and designed the study, supervised the work, and prepared the manuscript.

## Supplementary figure legends

**Fig. S1: Dorsal forebrain organoids to study the effects of different antiepileptic drugs (AEDs).**

**A.** Schematic of dorsal forebrain organoid generation(23) and AED treatment from day 21-35. Ctrl, no treatment (purple); VPA, Valproate treatment (orange); and LEV, Levetiracetam treatment (green). Three individual DFO batches were generated using two different iPSC cell lines (BIONi010-C and HMGU1).

**B.** Representative light microscope images of organoid development during treatment period for Ctrl, 1 mM VPA and 1mM LEV.

**C.** Quantification of organoid size throughout treatment period with VPA compared to Ctrl. Data points represent the mean of all data in each condition, and the error bars represent the standard error of mean (SEM). For all concentrations of VPA and LEV at D20, D24, D30 and D35, n=14 (except for 1 mM VPA and 0.5 mM VPA at D35 n=13; 0.5 mM LEV at D20 n=13) and at D27 n=10; for Ctrl at D20 and D35 n=29, at D24 n=30, D27 n= 22 and at D30 n=28. One-way ANOVA test with Bonferroni correction for multiple comparisons was used for significance testing between VPA and Ctrl per time point and no significance was found for any treatment to Ctrl comparisons.

**D.** 50 μm-wide cropped confocal image of a representative VZ-like region. RG cells (magenta) and proliferative cells (cyan).

**E.** Quantification of Ki67+SOX2+ over total SOX2+ cells in this region for each treatment condition (Ctrl, n=26; 0.1 mM VPA, n=28; 0.5 mM VPA, n=24; 1 mM VPA, n=26; 0.1 mM LEV, n=22; 0.5 mM LEV, n=16; 1 mM LEV, n=19). One-way ANOVA statistical test was performed with Bonferroni correction for multiple testing, only comparison between Ctrl vs. 1 mM VPA was significant (p=0.0039).

**F.** Representative image of DFO section for SOX2+ RG cells (magenta) and cCas3+ apoptotic cells (cyan).

**G.** Quantification of apoptosis in vRG cells in 50 μm-wide ventricular area in DFO slice at D35. Nonparametric Mann-Whitney test was performed for statistical analysis and showed no significance between Ctrl (n=22) and 1 mM VPA (n=24) groups. Shapiro-Wilk test was used to assess normality.

**Fig. S2: Dorsal forebrain organoids to study the effects of long-term exposure to VPA.**

**A.** 50 μm-wide cropped image of VZ-like region for the quantification of Ki67+SOX2+/Total SOX2+ cells. An example of double positive Ki67+SOX2+ cell is shown using a white arrow.

**B.** Immunohistochemistry staining of Vimentin (cyan) and SOX2+ RG cells (magenta). Three individual DFO batches were generated using two different iPSC cell lines (BIONi010-C and HMGU1).

**C.** Quantification of Vimentin signal at the organoid edges. Welch’s t-test was performed for statistical test (p=0.0013, n=8 for Ctrl and n=9 for VPA).

**Fig. S3: WGCNA network analysis with RNA-sequencing data.**

**A.** Dendrogram for the WGCNA to identify the cluster of genes with similar expression patterns.

**B.** Selected GO terms for biological processes associated with upregulated genes in the WGCNA yellow module.

**C.** Selected GO terms for biological processes associated with downregulated genes in the WGCNA brown module.

**D.** Selected GO terms for biological processes associated with downregulated genes in the WGCNA black module.

**Fig. S4: Quality control measurements for scRNAseq**

**A.** Violin plot of QC metrics, including number of features, number of gene counts, ribosomal gene counts and mitochondrial gene counts after filtering.

**B.** UMAP plot with crude cell type annotations after reciprocal rPCA integration of the data.

**C.** Stressed cell assessment by Gruffi. Distribution of the stressed (cyan) and not stressed (red) cells on UMAP (top left), distribution of conditions on UMAP (top right) and proportion of stressed cells within each cell types (bottom).

**D.** UMAP plot of normalized cells after integration by sample, with annotations for new cluster numbers (top left), cell cycle phase (top right), distribution of cells from different samples (bottom left), and distribution of cells from different cell lines (BIONi010-C and KOLF2.1J) (bottom right), following the removal of off-target and stressed cells.

**E.** UMAP plots showing technical features, including number of RNA features (top left), RNA count (top right), ribosomal RNA counts (bottom left), and mitochondrial RNA counts (bottom right) after removal of stressed cells.

**Fig. S5: Cell type annotation and cell-type specific analysis for scRNAseq**

**A.** Heatmap of manually selected cell type markers used for cell type annotation.

**B.** UMAP plot showing cell type annotations based on reference-query integration with a neocortical development dataset (left) and the respective confidence scores (right). As reference dataset, integrated from multiple different fetal transcriptomic datasets were used(52).

**C.** Stacked bar plot with the proportion of cell types per sample. All samples from different cell lines and conditions were plotted individually.

**D.** Volcano plot of DEGs in IPC/Newborn ExN cells comparing transcriptome of VPA-treated DFOs to Ctrl (significance threshold: absolute log2 Fold change > 1.5 and FDR < 0.0001). Each datapoint represents a gene; upregulated DEGs are colored in purple, downregulated DEGs are in blue and nonsignificant genes are in grey.

**E.** Bar plot showing selected biological processes derived from GO analysis using significantly upregulated DEGs between IPC/Newborn ExN cells from VPA and Ctrl DFOs.

**F.** CellChat analysis of cell-cell communication comparing VPA and Ctrl transcriptomes. Bar plot of interaction strength (left) and relative information flow (right). VPA in cyan, Ctrl in red.

**Fig. S6: Transcriptomic analysis of VPA-treated and Ctrl DFOs**

**A.** Heatmap showing the mean AUC scores of the top 50 most active regulons in RG cells as assessed with SCENIC, and across experimental conditions, VPA and Ctrl.

**B.** Heatmap displaying the enrichment of ASD-related gene sets in VPA-treated cell types. The enrichment is calculated based on the log2 odds ratio from Fisher’s exact test, with p-values annotated on each tile. Significant enrichment is shown across multiple cell types and gene sets, filtered by FDR < 0.0001 and log2FC > 1.5. Gene sets are categorized as SFARI 2019, 2022, and 2024. Colors represent the magnitude of enrichment (log2 odds ratio).

**Fig. S7: Proteome analysis of DFOs treated with VPA compared to Ctrl**

**A.** PCA plot of all VPA-treated (orange) and Ctrl (purple) proteome samples from D20, D35 and D50. Each datapoint represents one sample, n=9 per time point. As shown in Fig. 5A, DFOs were treated with 1 mM VPA from D21 until D50. BIONi010-C, KOLF2.1J and HMGU1 cell lines were used.

**B.** Volcano plot of differentially abundant proteins with increased abundance (purple) and decreased abundance (blue) proteins comparing VPA vs. Ctrl DFO samples at D35.

**C.** Volcano plot of differentially abundant proteins with increased abundance (purple) and decreased abundance (blue) proteins comparing VPA vs. Ctrl DFO samples at D50.

**D.** Bar plot showing the number of differentially abundant proteins for VPA vs. Ctrl DFOs at D35 and D50. For **B**, **C** and **D** the difference between time points is plotted against -log(P-value) with significance threshold: log2 Fold change value ≥1, FDR<0.05).

