## Supplementary figures and images for "Multiomics analysis identifies VPA-induced changes in neural progenitor cells, ventricular-like regions, and cellular microenvironment in dorsal forebrain organoids"

### Supplementary figure 1

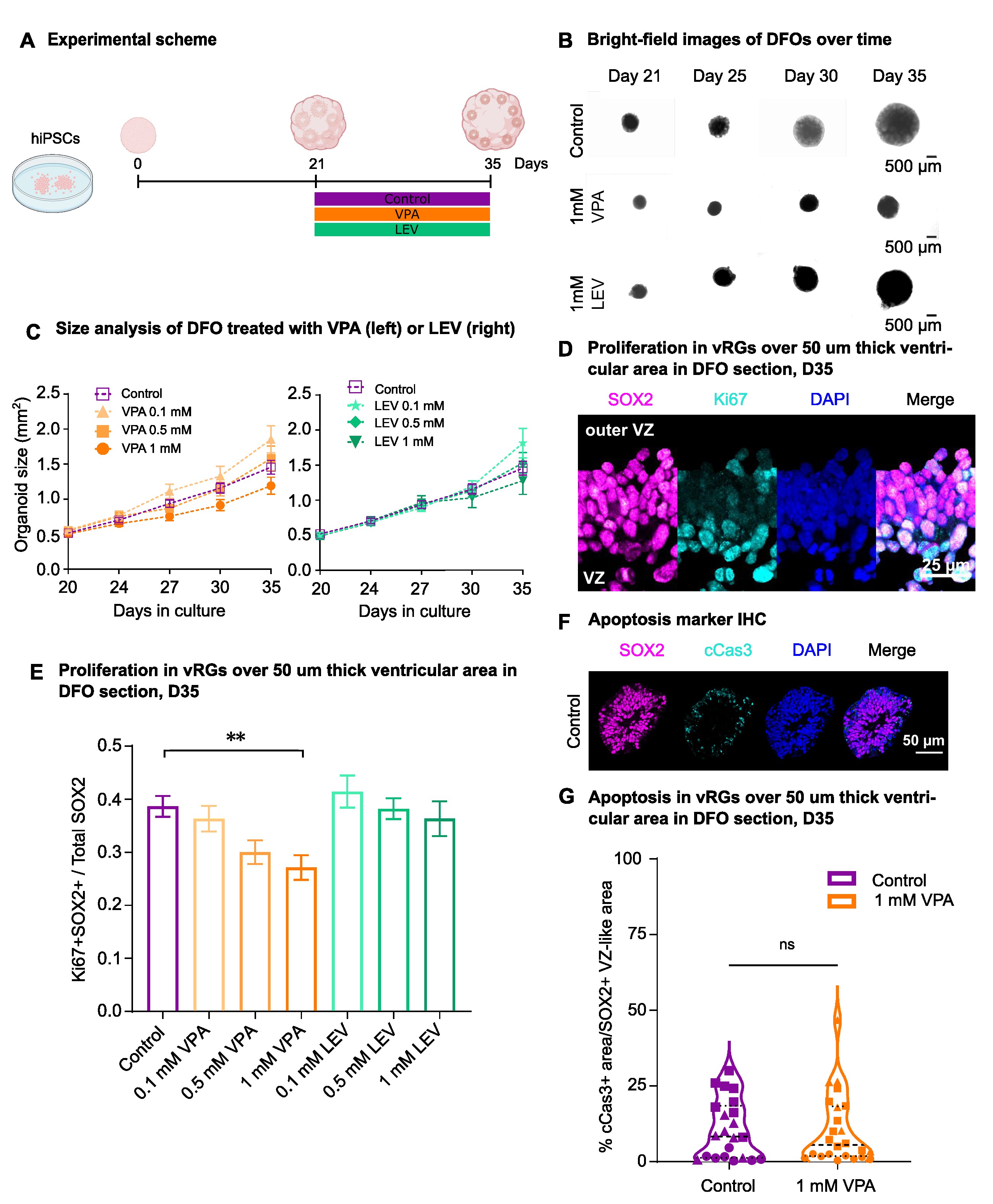

### Supplementary figure 2

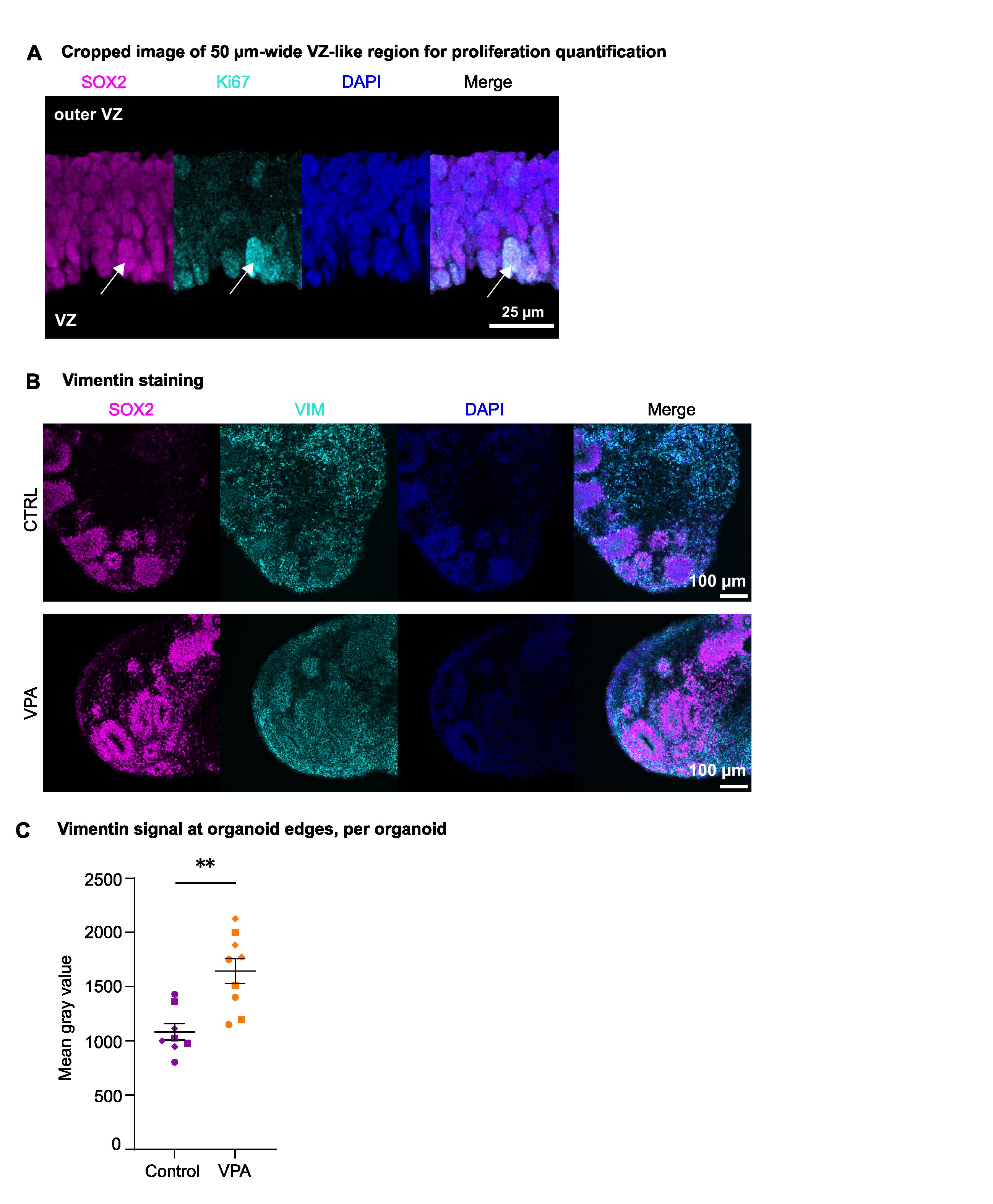

### Supplementary figure 3

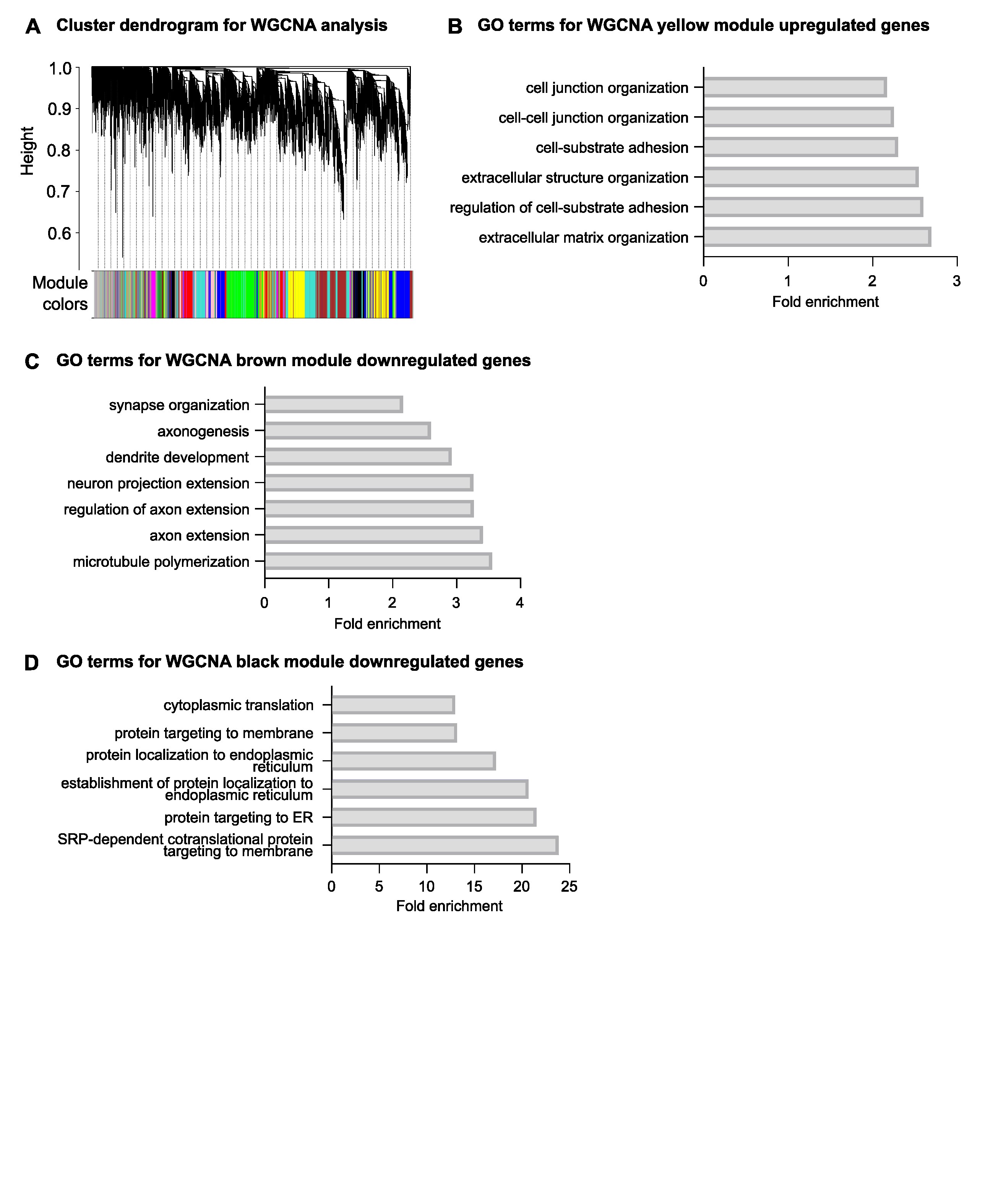

### Supplementary figure 4

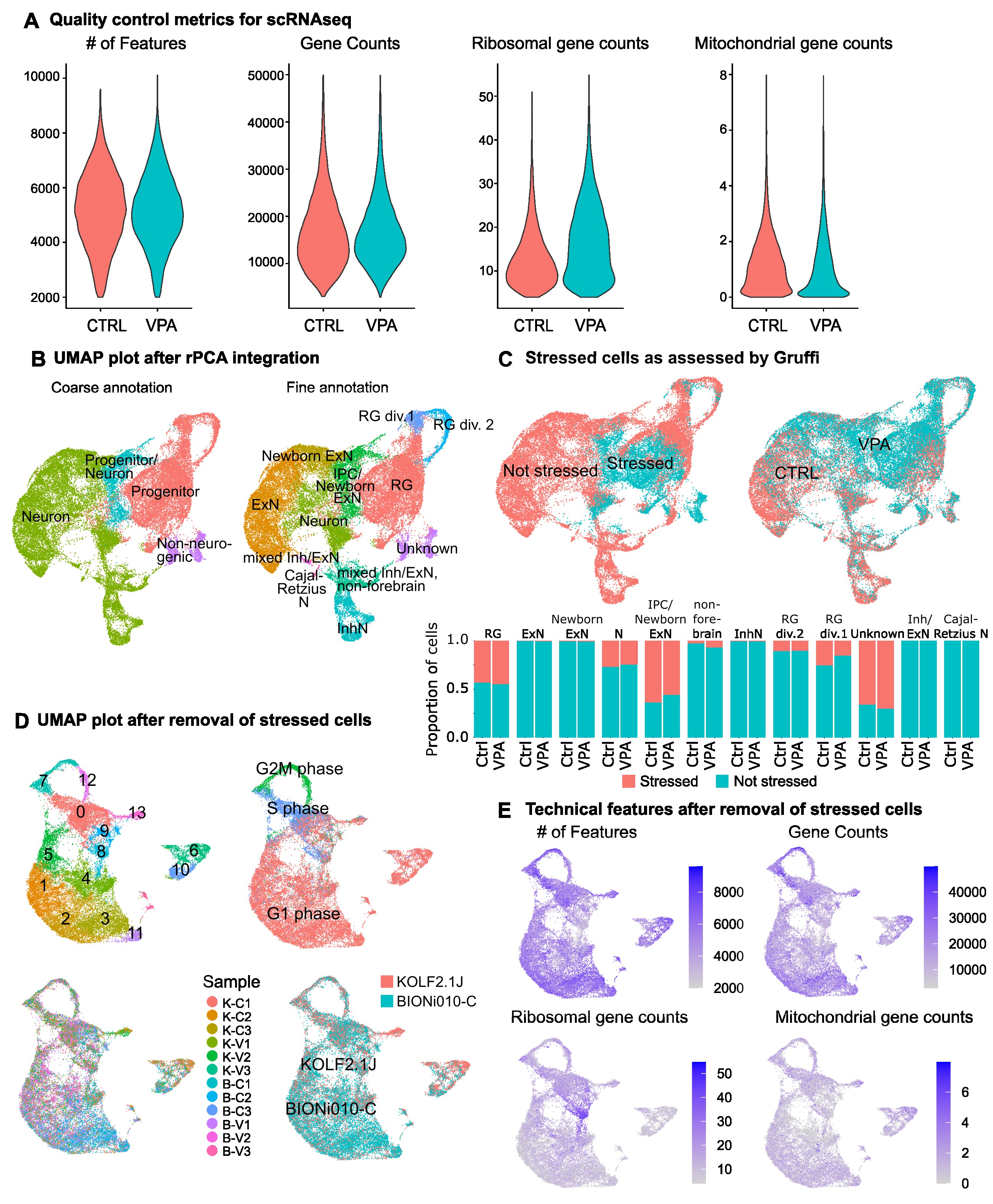

### Supplementary figure 5

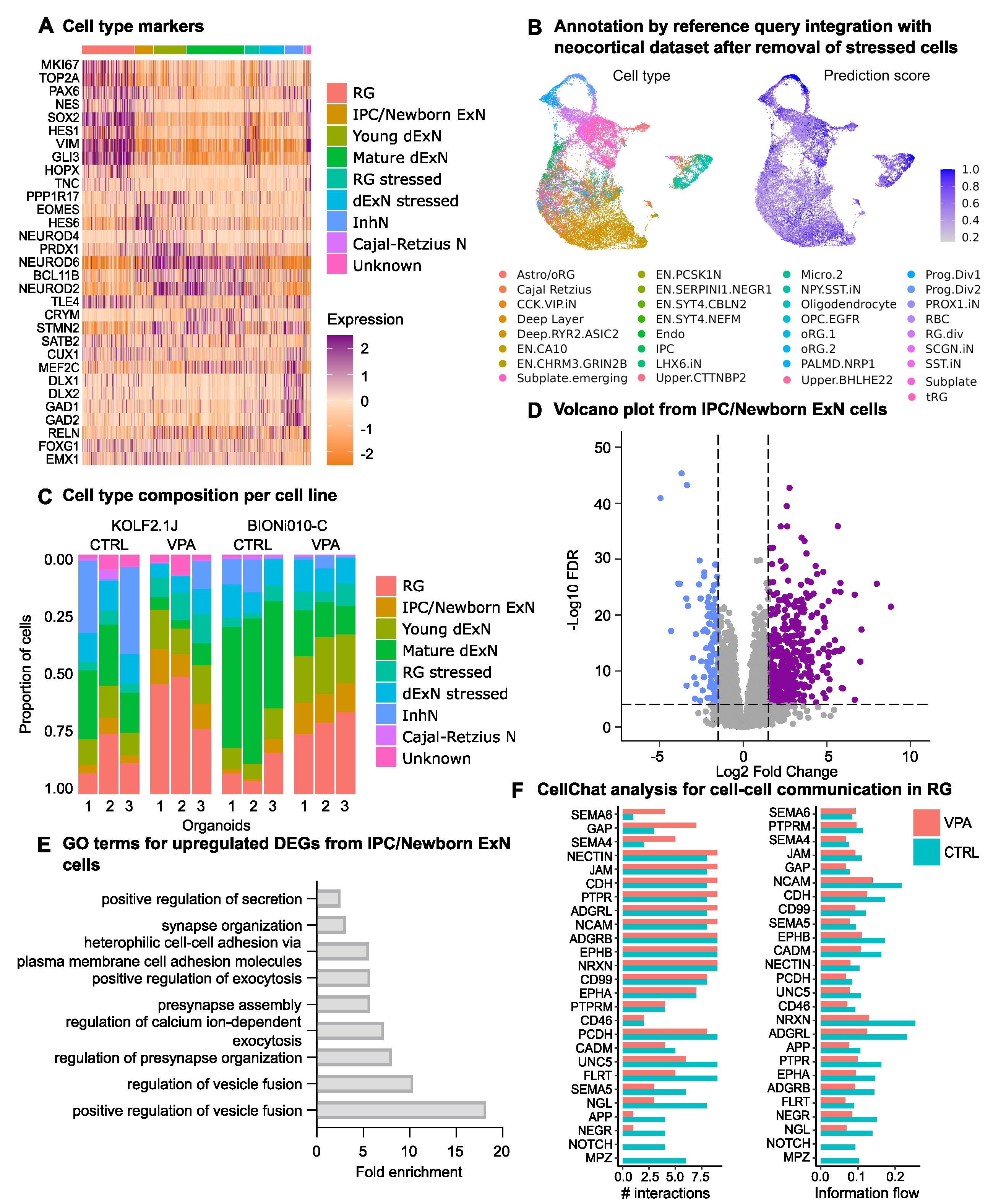

### Supplementary figure 6

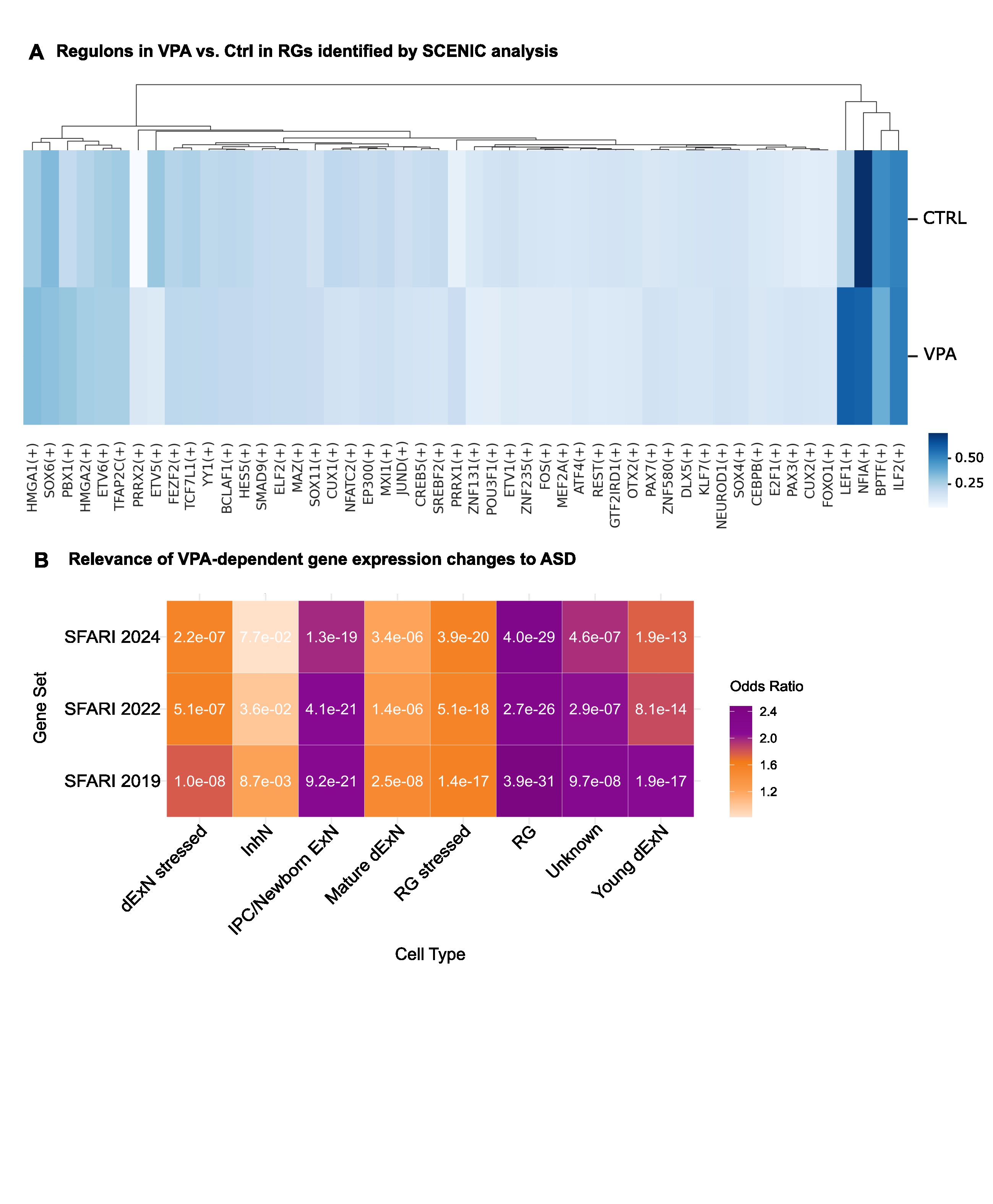

### Supplementary figure 7

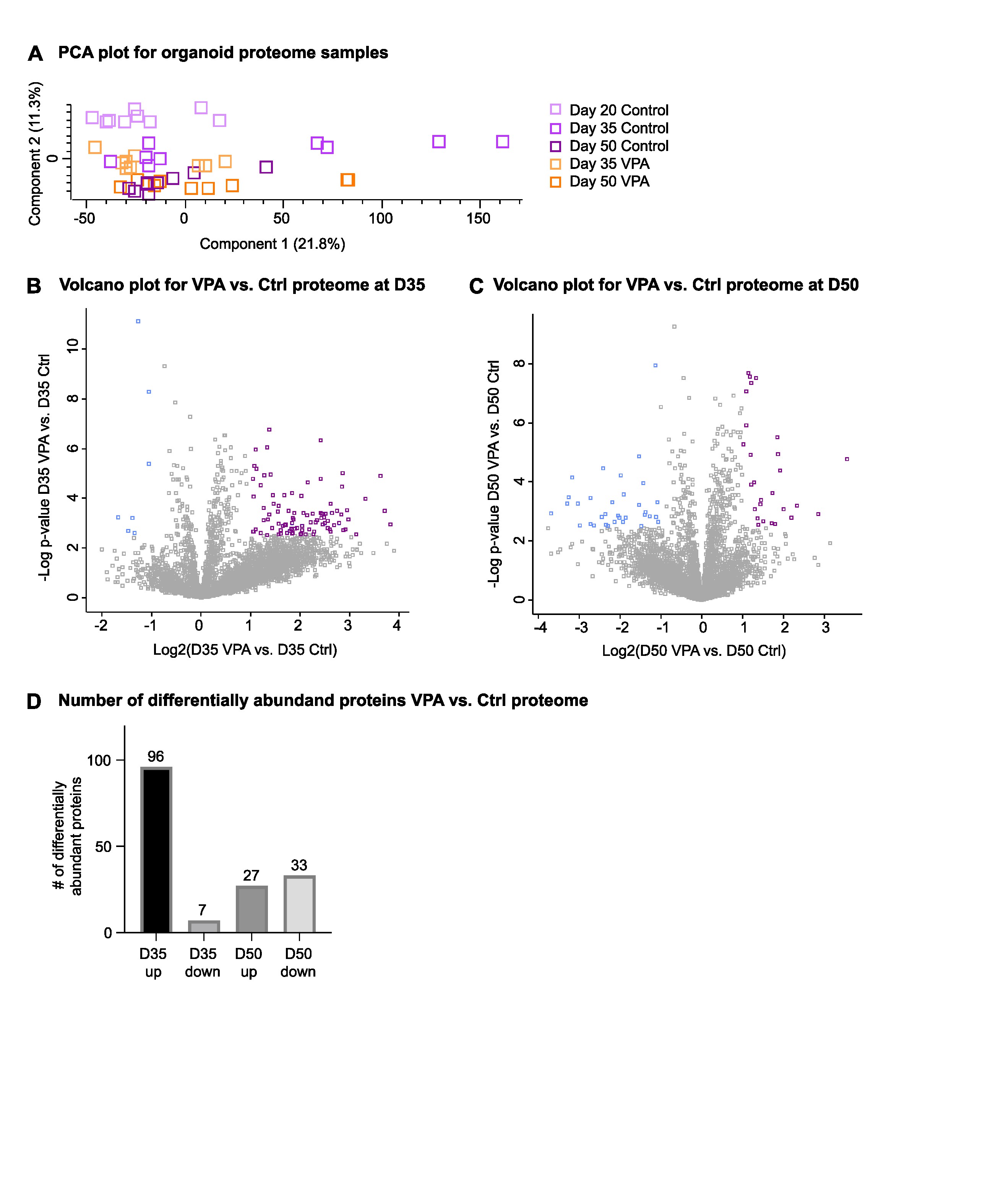
